# The population genomics of increased virulence and antibiotic resistance in human commensal *Escherichia coli* over 30 years in France

**DOI:** 10.1101/2021.06.24.449745

**Authors:** Julie Marin, Olivier Clermont, Guilhem Royer, Mélanie Mercier-Darty, Jean Winoc Decousser, Olivier Tenaillon, Erick Denamur, François Blanquart

**Affiliations:** Université Sorbonne Paris Nord, INSERM, IAME, 93017 Bobigny, France; Université Paris Cité, INSERM, IAME, 75018 Paris, France; Génomique Métabolique, Genoscope, Institut François Jacob, CEA, CNRS, Univ Evry, Université Paris-Saclay, 91042 Evry, France; AP-HP, Département de prévention, diagnostic et traitement des infections, Hôpital Henri Mondor, 94000 Créteil, France; AP-HP, Next Generation Sequencing Platform, University Hospital Henri Mondor, 94000 Créteil, France; EA 7380 Dynamyc Univ Paris Est Creteil (UPEC), Ecole Nationale Vétérinaire d’Alfort (EnvA), Faculté de Médecine de Créteil, 94000 Créteil, France; AP-HP, Laboratoire de Génétique Moléculaire, Hôpital Bichat, 75018 Paris, France; Center for Interdisciplinary Research in Biology, CNRS, Collège de France, PSL Research University, 75005 Paris, France

**Author notes:** co-last authors (Erick Denamur, François Blanquart).

**Keywords:** antibiotic resistance, commensal, *Escherichia coli*, evolution, genomics, virulence

## Abstract

*Escherichia coli* is a commensal species of the lower intestine, but also a major pathogen causing intestinal and extra-intestinal infections, increasingly prevalent and resistant to antibiotics. Most studies on genomic evolution of *E. coli* used isolates from infections. Here instead, we whole-genome sequenced a collection of 403 commensal *E. coli* isolated from fecal samples of healthy adult volunteers in France (1980-2010). These isolates were distributed mainly in phylogroups A and B2 (30% each) and belonged to 152 sequence types (STs), the five most frequent being ST10 (phylogroup A) (16.3%), ST73 and ST95 (phylogroup B2) (6.3 and 5.0%, respectively), ST69 (phylogroup D) (4.2%) and ST59 (phylogroup F) (3.9%), and 224 O:H serotypes. ST and serotype diversity increased over time. The O1, O2, O6 and O25-groups used in bioconjugate O-antigen vaccine against extra-intestinal infections were found in 23% of the strains of our collection. The increase in frequency of virulence-associated genes and antibiotic resistances was driven by two evolutionary mechanisms. Evolution of virulence gene frequency was driven by both clonal expansion of STs with more virulence genes (“ST-driven”) and increases in gene frequency within STs independently of changes in ST frequencies (“gene-driven”). In contrast, the evolution of resistance was dominated by increases in frequency within STs (“gene-driven”). This study provides a unique picture of the phylogenomic evolution of *E. coli* in its human commensal habitat over 30 years and will have implications for the development of preventive strategies.

**IMPORTANCE:** *Escherichia coli* is an opportunistic pathogen with the greatest burden of antibiotic resistance, one of the main causes of bacterial infections and an increasing concern in an ageing population. Deciphering the evolutionary dynamics of virulence and antibiotic resistance in commensal *E. coli* is important to understand adaptation and anticipate future changes. The gut of vertebrates is the primary habitat of *E. coli* and probably where selection for virulence and resistance take place. Unfortunately, most whole-genome sequenced strains are isolated from pathogenic conditions. Here, we whole genome sequenced 403 *E. coli* commensals isolated from healthy French subjects on a 30-year period. Virulence genes increased in frequency by both clonal expansion of clones carrying them and increases in frequency within clones whereas resistance genes increased by within clone increased frequency. Prospective studies of *E. coli* commensals should be performed worldwide to have a broader picture of evolution and adaptation of this species.

## INTRODUCTION

*Escherichia coli* is a commensal species of the lower intestine of humans and other vertebrates (1). It is also a major opportunistic pathogen causing intestinal and extra-intestinal infections (2), ranked third in the World Health Organization list of antibiotic resistant ‘priority pathogens’, as it is today the bacterial pathogen associated with the largest burden of antimicrobial resistance (3).

*E. coli* strains share a core genome of about 2,000 genes over 4,700 genes in average (4). In spite of numerous recombination events, this species presents a robust clonal structure with at least nine phylogroups called A, B1, B2, C, D, E, F, G and H (2), and sequence types (STs) defined at a finer genetic resolution. In addition, molecular typing of genes coding for surface structures as O-polysaccharide and H-flagellar antigens (5), as well as the FimH protein (6), allows, in conjunction with STs, a better characterization of the clones (7). O-antigens are key elements in the pathophysiological processes and used as a target for the development of *E. coli* vaccines to protect against extra-intestinal infections (8, 9).

From a public health perspective, it is particularly important to understand the evolution of the repertoire of virulence-associated and antibiotic resistance genes. Virulence genes are associated with a higher risk of urinary tract and bloodstream infections (10, 11). Antibiotic resistant infections are associated with longer hospital stay, increased risk of death and public health costs (12). Both virulence and antibiotic resistance genes repertoire greatly varies among strains, with continuous gene acquisition and loss (13). Temporal trends in virulence have been less often examined than trends in resistance. The mean number of virulence genes in commensal *E. coli* has increased from 1980 to 2010 in France (14). Increased virulence of *E. coli* could be one explanation for the increasing incidence of *E. coli* bacteremia in Europe (12, 15), together with increased reporting or changing epidemiological factors like the ageing population. Resistances to multiple antibiotics have concomitantly rapidly increased in frequency in *E. coli* over the last decades, and stabilized at an intermediate level (16).

What evolutionary forces act on virulence and resistance genes? A first hypothesis is that these genes are direct targets of selection. Virulence genes are thought to primarily be selected in the intestine as a by-product of commensalism (17, 18). Virulence genes are linked with longer persistence in the gut (19). Extra-intestinal compartments do not typically lead to efficient onward transmission and are often considered an evolutionary dead end (18). Alternatively, clonal expansion could modify the prevalence of resistance and virulence genes, without these genes being the direct target of selection. Clonal expansion may be important in *E. coli* since several resistance and virulence genes are closely linked to specific STs (20, 21). The selective forces acting on resistance are clearer. Resistance is selected by antibiotic use, as evidenced by the spatial correlation between local antibiotic use and resistance (22), and the fact that hosts are colonized by resistant strains more frequently after antibiotic use (23). For *E. coli*, exposure to antibiotics results 95% of the time from antibiotic treatment prescribed for infections unrelated to *E. coli*, and not to treat an *E. coli* infection (24). Such “bystander” antibiotic exposure happens around once per year in adults in France (25).

Although the gut is the typical habitat of *E. coli* and likely the main ecological context of selection for virulence and resistance, most studies on the evolution of virulence and antibiotic resistance focused on pathogenic clinical strains isolated from extra-intestinal infections (26–28). Extra-intestinal infections are a rare occurrence in *E. coli*’s life: the incidence of urinary tract infections is of the order of 10^-2^ per person-year (29), and of bacteremia of the order of 10^-4^ per person-year (30). In comparison to commensal *E. coli*, *E. coli* sampled from infections are biased towards strains carrying more virulent genes (11, 26, 27). Strains from infections may also be less diverse than commensal strains as they belong to some specific clones (2, 31), hindering the detection of temporal trends in these elements and the evolutionary mechanisms driving the emergence of pathogenic clones (32).

To avoid these biases, we investigated here the short-term genomic evolution of commensal *E. coli* over 1980 to 2010 using a rare collection of 403 strains from fecal samples of healthy volunteers in France. We assembled this collection from five previously described collections all collected in healthy individuals with a diversity of ages, sex, and located in two French regions (table S1), and controlled for these covariates in all analyses.

## RESULTS

The pan-genome of the 403 strains contained 47,367 genes, with no saturation in the number of genes detected (figure S1). The core genome, defined as the number of genes found in more than 99% of the isolates, contained 2156 genes. A total of 504 genes were found in 95% to 99% of the isolates and 3252 genes were found in 15% to 95% of the isolates. The large majority of genes (41,455 out of 47,367) were found in less than 15% of the isolates.

### 1. Evolution of phylogenetic groups, STs and serotypes over time

Of the 403 isolates, 398 were assigned to the *E. coli* phylogroups A (120, 30%), B1 (58, 14%), B2 (120, 30%), C (10, 2%), D (44, 11%), E (13, 3%), F (26, 6%), G (6, 1%) and H (1, 0.2%) (tables S2-3). We found two strains (0.5%) belonging to the *Escherichia* clade IV and one (0.2%) to the *Escherichia* clade V. Two strains (C020-013 and LBC20a, 0.5%) were phylogenetically distinct from the known phylogroups (A to H). They probably belong to an undescribed phylogroup (figure S2). We fitted a binomial generalized linear model (GLM) for the presence/absence of each phylogroup as a function of time, controlling for host age, region, sex. The use of antibiotics in the past month was an exclusion criterion in four of the five studies generating the collections. For the last collection, we controlled for the possible (but unlikely) use of antibiotic by adding a specific factor for this collection in the regression (table S1). We detected a significant decrease in frequency for the phylogroup A (from 58% in 1980 to 26% in 2010, p-value=1.79e-03) and a significant increase in frequency for the phylogroup B2 (from 9% in 1980 to 36% in 2010, p-value=1.34e-03) (figure 1A). No significant change was detected for the other phylogroups.

**Figure 1.**
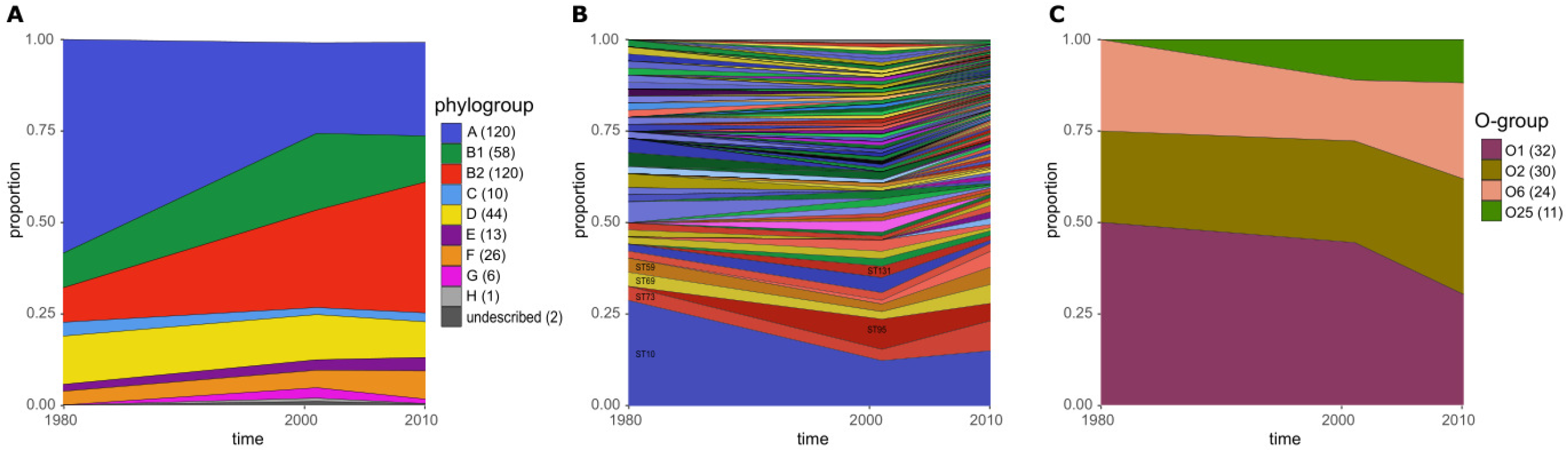
Frequency distribution of phylogroups and STs between 1980 and 2010. (A) Frequency distribution of phylogroups. The sample size is indicated in brackets. The decline of phylogroup A and the increase of phylogroup B2 were significant (0.05 level). (B) Frequency distribution of STs. The proportion of each ST has been plotted by year ordered by the overall frequency (most common at the bottom). Only the names of STs with an overall frequency > 0.03 and ST131 are shown. The only variation that was significant at the 0.05 level was the decline of ST10. (C) Frequency of the O-groups O1, O2, O6 and O25. The sample size is indicated in brackets. No significant variation with time was detected (0.05 level).

Those 403 isolates were distributed among 152 Warwick University scheme STs (33). The five most frequent STs were ST10 (15.4%) (phylogroup A), followed by ST73 (5.9%) and ST95 (4.7%) (both of phylogroup B2), ST69 (4.0%) (phylogroup D) and ST59 (3.7%) (phylogroup F) (figure 1B and table 1). ST131 represented only a small fraction of our collection (1.6%) and increased in frequency from 0.0% in 1980 to 1.3% in 2010. Using a binomial GLM, we detected a significant decrease in frequency for ST10 between 1980 and 2010 (from 29% in 1980 to 15% in 2010, p-value=0.028). No other significant change was detected for the three others most frequent STs. A total of 30, 59 and 98 distinct STs were detected in 1980, 2001 (2000–2002) and 2010 respectively. ST diversity increased over time even after accounting for the increasing number of strains sampled, and the changes in phylogroup frequencies (figures 1B and S3).

**Table 1.** Distribution of the 11 most frequent sequence types of the *E. coli* commensal collection isolates. The number of isolates and the percentage for each year are presented in the table (see table S12 for the complete table). The year 2001 corresponds to the aggregated data of years 2000, 2001 and 2002. The * indicate the sequence types (STs) for which we inferred the divergence times.

| ST (phylogroup) | Commensal |  |  |  |
| --- | --- | --- | --- | --- |
|  | 1980 | 2001 | 2010 | All years |
| 10* (A) | 15 (29%) | 12 (12%) | 35 (15%) | 62 (16%) |
| 73 (B2) | 2 (4%) | 3 (3%) | 19 (8%) | 24 (6%) |
| 95* (B2) | 0 (0%) | 8 (8%) | 11 (5%) | 19 (5%) |
| 69* (D) | 2 (4%) | 2 (2%) | 12 (5%) | 16 (4%) |
| 59 (F) | 2 (4%) | 2 (2%) | 11 (5%) | 15 (4%) |
| 141 (B2) | 0 (0%) | 1 (1%) | 10 (4%) | 11 (3%) |
| 93 (A) | 1 (2%) | 4 (4%) | 2 (1%) | 7 (2%) |
| 452 (B2) | 1 (2%) | 2 (2%) | 5 (2%) | 8 (2%) |
| 405 (D) | 1 (2%) | 2 (2%) | 2 (1%) | 5 (1%) |
| 58 (B1) | 0 (0%) | 2 (2%) | 3 (1%) | 5 (1%) |
| 131 (B2) | 0 (0%) | 3 (3%) | 3 (1%) | 6 (2%) |
| **Total in collections** | **53** | **104** | **246** | **403** |

We next typed the isolates using the diversity of genes coding for O and H antigens and FimH protein. These genes are located in the two major hot spots of recombination along the *E. coli* chromosome and the corresponding antigens are important for the host immune response and pathogenesis (4). Serotype (O:H types genetically defined) diversity increased between 1980 and 2010, however, the diversity of O-groups, H-types and *fimH* alleles remained stable between 2001 and 2010 when considered individually (figure S4 and tables S4-7). The four O-groups targeted by the recently developed bioconjugate vaccine ExPEC4V (8, 9), O1, O2, O6 and O25 represented 24% of our collection. By fitting a binomial generalized linear model (GLM), we did not detect a change in frequency between 1980 and 2010 (figure 1C).

In the analyses of temporal trends in phylogenetic groups, STs, and serotypes over time, we controlled for the covariates host age, regions, sex and possible use of antibiotics. In this series of linear models, only 3 out of 88 inferred effects of control covariates were significant at the 0.05 level. This suggested that these covariates have no or negligible effect on the genetic composition of *E. coli*.

We next examined how the diversity of O and H antigen and FimH protein coding genes evolves in relation to the phylogenetic structure of three STs with a molecular clock signal and that we were thus able to date: ST10, ST69 and ST95. ST10 strains exhibited a large diversity of O and H antigens (38 serotypes) and *fimH* alleles (15 alleles). ST69 strains exhibited a limited diversity of O serogroups (O15, O17, O25 and O45) associated with H18 and *fimH*27 almost exclusively (figure 2). We found six serotypes and five *fimH* alleles among ST95 strains. The apparent diversity in O:H combinations and *fimH* alleles reflected in part the age of the ST that can vary by a 20-fold factor according to our estimations (figure 2). ST10 started to diversify hundreds of years before ST69. The MRCA (most recent common ancestor) dates of ST10, ST69 and ST95 were 1377 [866-1637], 1951 [1924-1969], and 1680 [766-1882] respectively (Bayesian skyline model, figure S5 and table S8). We tested whether the difference in their timescale of evolution explains the difference in diversity and found that the diversity of genes encoding for O and H antigens and for the FimH protein increased faster for the more recent ST69 than for ST10 and ST95 (figure S6). We also quantified the change in diversity of O:H serotypes and *fimH* alleles within the most abundant ST of our commensal collection (ST10). We did not detect a significant change in frequency for any of the five most frequent serotypes within ST10 between 1980 and 2010 (figure S7).

**Figure 2.**
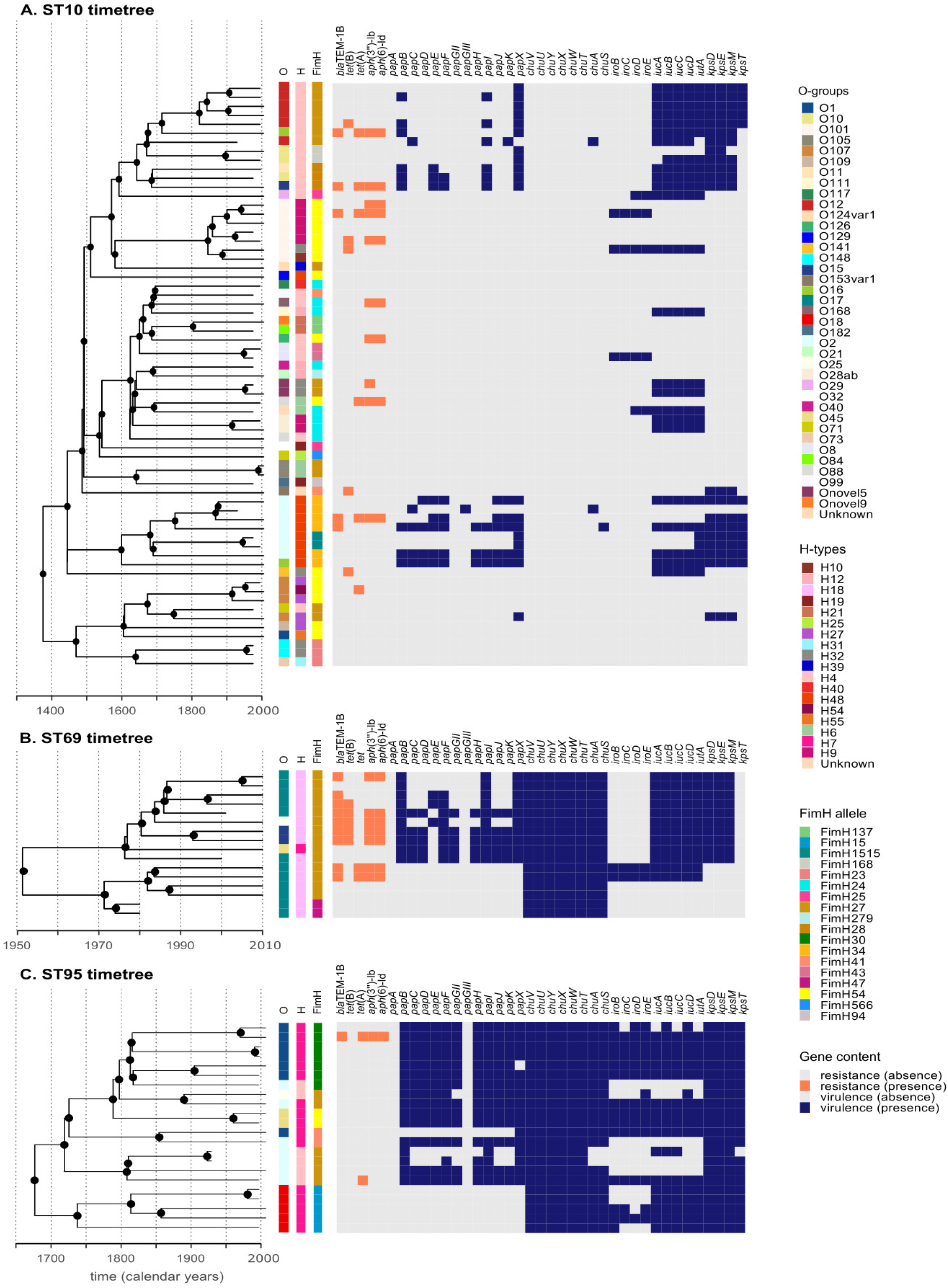
Genomic content of three of the most frequent STs. For ST10 (A), ST69 (B) and ST95 (C), in addition to the diversity of O:H serotypes and *fimH* alleles, we examined the presence of five antibiotic resistance genes (*bla*_TEM-1B_, *tet*(B), *tet*(A), *aph*(3’’)-Ib and *aph*(6)-Id) and of 34 virulence genes [*pap* (adhesin), *chu*, *iro*, *iuc* (iron capture systems) and *kps* (protectin) operons]. The timetrees were built with BEAST v1.10.4 (65). ST10 exhibits the largest serotype diversity (44 serotypes), among the other STs examined here, ST69 (4 serotypes) and ST95 (6 serotypes). Nodes with a support value (Bayesian posterior probability) > 0.75 are indicated by black circles.

### 2. Recent temporal evolution of virulence and resistance genes in commensal *E. coli*

We first evaluated the effect of phylogroup frequency changes and time on the recent temporal evolution of virulence and resistance genes. Next, we analyzed the gene frequency variations over two periods of time, between 1980 and 2001 and between 2001 and 2010. We compared the change in frequency of virulence and antibiotic resistance genes with that of non-resistance and non-virulence (NVNR) genes over each time period. This tests whether virulence and resistance genes change in frequency significantly faster than NVNR genes, which may reflect the action of selection. We then quantified potential directional selection on gene presence with a logistic model (gene presence/absence *versus* time as a continuous variable) between 1980 and 2010. Finally, we computed dN/dS ratios to test if gene sequences evolved neutrally or were under diversifying or purifying selection.

#### a. Recent increase in virulence and resistance genes

The correspondence analysis indicated a significant association between the phylogroups and the virulence and resistance gene categories (chi-squared test, p-value < 2.2e-16) (figure S8 and tables S9-11). The phylogroup B2 carried more virulence factors (VFs) distributed in the adhesin, invasin, iron acquisition and protectin categories, but fewer resistance genes. The phylogroups A, B1 and E carried more toxin genes. The phylogroups C, D and the undescribed phylogroup were positively correlated with the large majority of resistance categories, including resistances to beta-lactam, sulphonamide, aminoglycoside, phenicol, tetracycline and trimethoprim antibiotics.

The prevalence of virulence genes increased between 1980 and 2010 (figure 3), as already reported for a large part of this collection (14). First, the mean virulence score (number of virulence factors out of the 101 assayed) increased from 14 to 19. Using a linear model with sampling dates and phylogroups as predictors, we showed that, between 1980 and 2010, the increase in virulence score is explained by the change in the phylogroup frequency; the effect of the sampling date is not significant (effect size = −1.90e-02 [-0.140; 0.102] VF product per year, p-value = 0.743). This increase was driven by the change in frequency of several phylogroups carrying many VFs, such as B2 (9% of B2 strains in 1980 to 36% in 2010 with a mean virulence score of 24, effect size = +11 [6; 17] compared to phylogroup A, p-value = 7.37-04) and F (4% of F strains in 1980 to 8% in 2010 with a mean virulence score of 23, effect size = +14 [7; 18] compared to phylogroup A, p-value = 1.87e-04). When testing each time period individually, we did not find a significant effect of the sampling date either, but only of the phylogroup frequency (between 1980 and 2001: effect size = +0.040 [-0.087; 0.167] VF product per year, p-value = 0.472; between 2001 and 2010: effect size = −0.325 VF product per year [-0.765; 0.114], p-value = 0.126).

**Figure 3.**
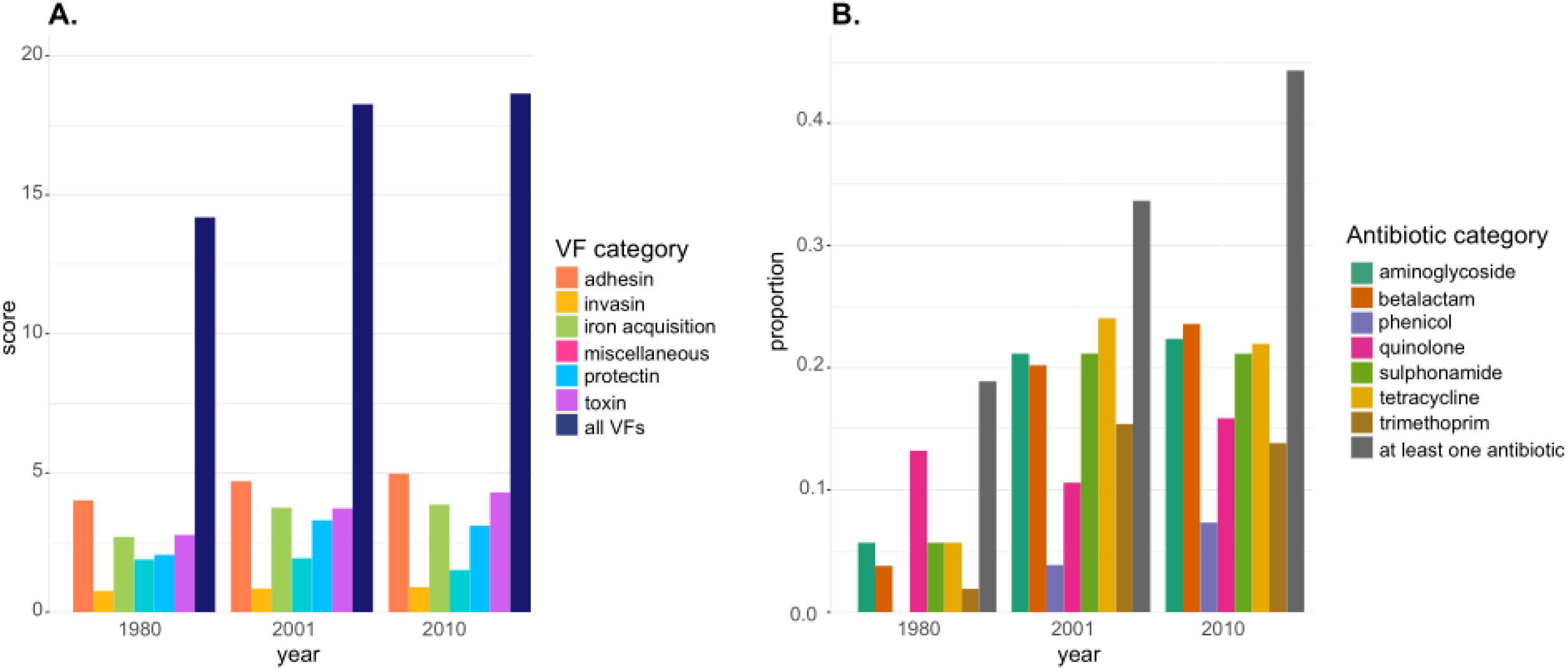
Evolution of virulence and antibiotic resistance genes between 1980 and 2010. (A) Mean virulence score computed for all the 403 commensal *E. coli* isolates. The virulence score of a strain is defined as the number of the VFs of each category tested that were present in that strain. (B) Frequency of antibiotic resistance through time for all 403 isolates. Both gene acquisition and point mutations are included, but we omitted the macrolide category because more than 99% of the strains were resistant to macrolide antibiotics. For readability, in each panel the year 2001 includes strains sampled in 2000, 2001 and 2002.

The frequency of resistance also increased through time (14). The fraction of strains resistant to at least one antibiotic class increased from 19% to 44% from 1980 to 2010 (figure 3). This rise was significant within a linear model with sampling dates and phylogroups as predictors (effect size = +1.12e-02 [4.47e-03; 1.79e-02] per year, p-value = 2.88e-03), showing increased resistance independently of changes in phylogroup frequencies. In addition to sampling dates, this increase was also driven by the change in frequency of one minor phylogroup, phylogroup F which increases from 4% in 1980 to 8% in 2010 and in which 88% of the strains are resistant to at least one antibiotic (effect size = +0.680 [0.361; 1.00] compared to phylogroup A, p-value = 3.87e-04). When considering each time period individually, the frequency of resistance to one antibiotic significantly increased between 1980 and 2001 (effect size = +9.77e-03 [4.13e-04; 0.019] per year, p-value = 0.043). The increase between 2001 to 2010 was comparable in size but not significant (effect size = +0.018 [-0.003; 0.039] per year, p-value = 0.082). Interestingly, the proportion of strains resistant to two or more antibiotic categories remained stable at 27% between 2001 and 2010.

We verified that genotypic resistance was a good predictor of phenotypic resistance (disk diffusion method) by comparing both approaches in a subset of 319 strains from the Coliville, PAR and VDG collections for the six following antibiotics: amoxicillin, nalidixic acid, cotrimoxazole, tetracycline, chloramphenicol and streptomycin. The genotype predicts the phenotype with good performance with an overall agreement of 95.1%, a sensitivity of 75.9%, a specificity of 98.4%, a positive predictive value of 89.2% and a negative predictive value of 95.9%.

#### b. ST clonal expansion versus increase in gene frequency within STs

We next evaluated how the temporal variation in the frequency of individual resistance and virulence genes was driven by clonal expansion or contraction of STs versus gene frequency changes within one or several STs. We chose to do this analysis at the level of STs because they are evolutionary units smaller than phylogroups but large enough to be followed through time, and distinct in their gene content (34). Considering the size of our collection, we would lack power to investigate temporal changes at a finer phylogenetic unit. Eight STs were present in 1980, 2001 and 2010, they belonged to the phylogroups A, B2, D and F and represent 35% of our dataset (table S12). We decomposed the temporal change in individual virulence and resistance gene frequencies in two additive components (figures 4-5 and tables S13-16). First, “ST-driven” change corresponded to a change in gene frequency driven by the change in frequency of STs (figure S9). Second, “gene-driven” change reflected the change in gene frequency within STs, independently of changes in ST frequencies.

**Figure 4.**
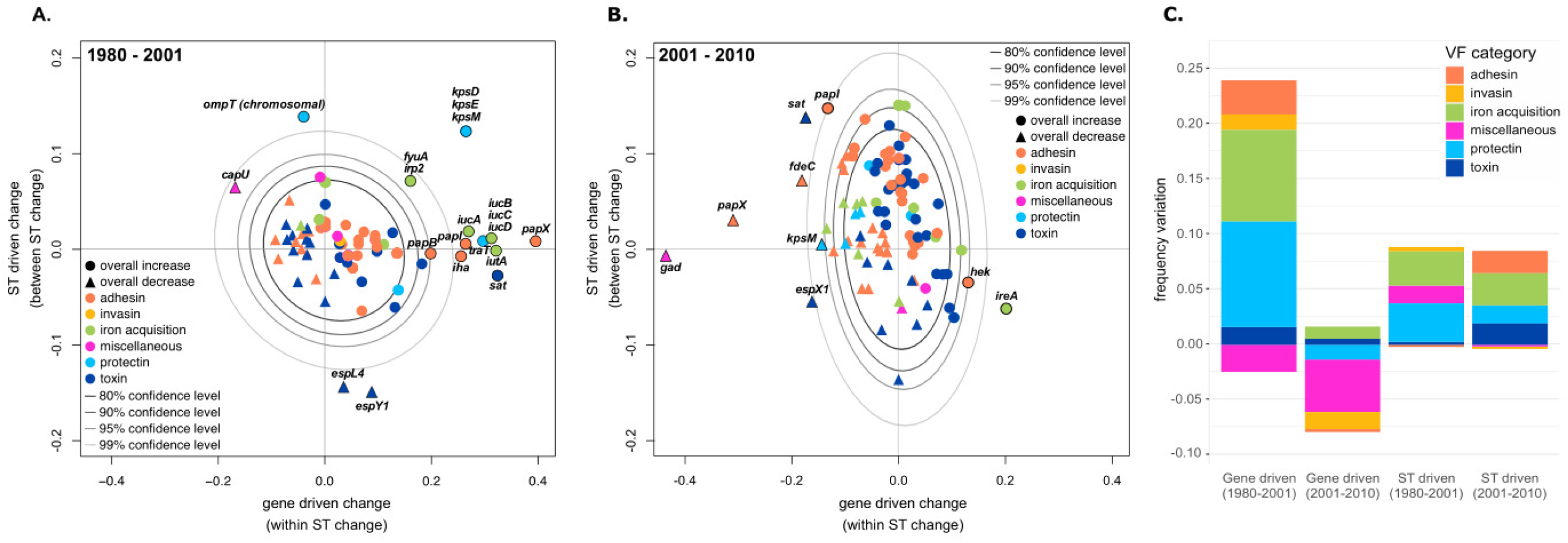
Temporal change of virulence gene frequency between 1980 – 2001 (A) and between 2001 – 2010 (B). The overall frequency change (*Δ*f**) for each gene (increases depicted by circles and decreases by squares) is decomposed in two additive components: change driven by the variation in frequency of STs carrying the focal gene (ST driven change) and change driven by the variation in frequency of the focal gene (gene driven change). For readability, only genes for which between ST change or within ST change was greater than 0.02 in absolute value are shown (see table S13-14 for the complete list). The grey ellipses represent the confidence levels computed for NVNR genes. Genes highlighted in bold are those for which the temporal change is significantly different from NVNR genes at the 0.05 level (out of the 95% confidence level). Summary of the two additive components of temporal change of virulence genes between 1980 and 2010 (C). We computed the mean change per product (e.g. mean change among *iucA*, *iucB*, *iucC*, *iucD* and *iutA* genes for aerobactin), and reported the mean change among products of a category (e.g. among products classified in the iron acquisition category).

**Figure 5.**
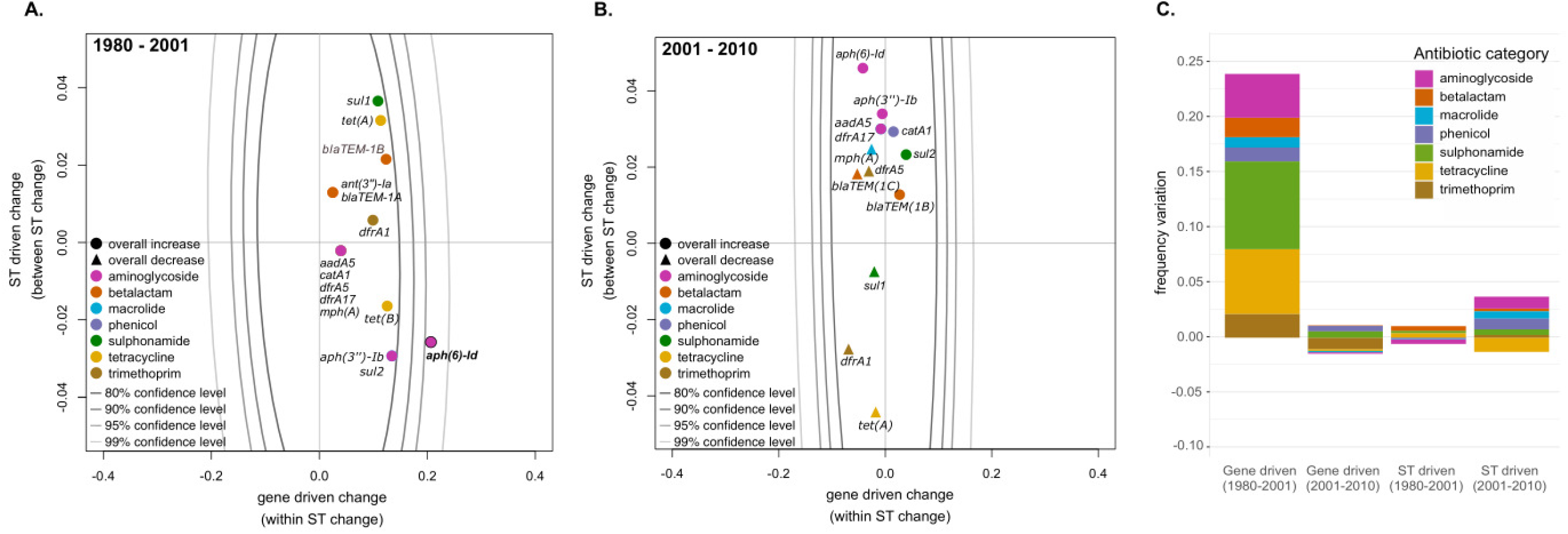
Temporal change of antibiotic resistance frequency between 1980 – 2001 (A) and between 2001 – 2010 (B). The overall frequency change (*Δ*f**) for each gene (increases depicted by circles and decreases by squares) is decomposed in two additive components, change driven by the variation in frequency of STs carrying the focal gene (ST driven change) and change driven by the variation in frequency of the focal gene (gene driven change). For readability, only genes for which between ST change or within ST change was greater than 0.02 are shown here (see table S15-16 for the complete list). The grey ellipses represent the confidence levels computed for NVNR genes. Genes highlighted in bold are those for which the temporal change is significantly different from NVNR genes at the 0.05 level (95% confidence level). Note the scale of the y-axis is much smaller for resistance than for virulence genes (figure 4) as ST-driven changes are minor compared to gene-driven changes for antibiotic resistance genes. Summary of the two additive components of the temporal change of antibiotic resistance genes between 1980 and 2010 (C). We computed the mean change among genes belonging to a given category (e.g. among *tet*(A), *tet*(B), *tet*(D) and *tet*(M) for tetracycline).

The evolution of virulence genes was both ST and gene driven. The relative contribution of the two processes depended on the time period considered. Frequency changes within STs (gene driven change) contributed mostly in the period between 1980 and 2001 (figure 4). Those changes, compared to change in frequency of NVNR genes, included notably the large increase in frequency of the *kps* operon (*kpsD*, *kpsE* and *kpsM*) involved in the K1 capsule biosynthesis (from 40% to 78%), of the *iuc* operon (aerobactin, from 32-36% to 60-68%), of the HPI (High Pathogenicity Island, from 52% to 75%) and of the *pap* operon (pyelonephritis-associated pilus, from 12-24% to 25-64%) (figure 4A and table S13). From 2001 to 2010, the change in virulence gene frequency was driven by both the increase in frequency of more virulent STs (ST driven change), in particular those carrying genes coding for adhesins, and the change in gene frequency within STs (gene driven change) (figure 4B). We retrieved similar results when removing phylogroup B2 strains, suggesting that the increase in frequency of virulence genes was not driven by the expansion of the B2 phylogroup (figure S10, tables S17-18).

In contrast to virulence genes, the evolution of resistance gene frequency was almost exclusively gene driven and occurred primarily between 1980 and 2001 (figure 5 and tables S15-16). When compared to change in frequency of NVNR genes, significant increase in frequency was detected for one gene responsible for resistance to aminoglycoside antibiotics (*aph*(6)-Id, from 0% to 18%). All resistance genes increased in frequency over that period. In contrast, resistance genes frequency did not change much in the second period between 2001 and 2010. We noted however that any increasing gene frequency was ST-driven. Thus, these genes rapidly increased in frequency in 20 years and subsequently stabilized, and their dynamics were unaffected by changes in the ST composition of the population. The same results were retrieved when removing B2 strains (figure S11, tables S19-20).

Gene content visualization of ST10, ST69 and ST95 corroborated these results (figure 2). Virulence genes appeared to be more phylogenetically clustered than resistance genes, explaining the importance of clonal expansion in driving the evolution of the virulence gene repertoire. We last tested if genes located on plasmids are more likely gene-driven (increasing simultaneously within multiple STs) than ST-driven (clonal expansion), but found no significant effect (correlation across genes between the proportion of gene driven change and plasmid *vs.* chromosome as a categorical variable, effect size = + 0.09 [-0.02; 0.21], p-value = 9.62e-02).

#### c. Further evidence for selection on virulence and resistance genes

We detected a significant increase in frequency over at least one period of time, by comparison with NVNR genes, for 25 virulence genes and one gene associated to antibiotic resistance (*aph*(6)-Id). To test if the presence of these genes (relative to absence) is under directional selection over the whole period 1980 to 2010, we fitted a logistic model for gene presence/absence as a function of time and phylogroup for these 26 genes. In population genetics, the action of selection on a gene results in a linear trend on the logit scale. Thus, the coefficient of a logistic model (binomial generalized linear model) is interpretable as a selection coefficient (additionally controlling for association with phylogroups). Out of the 17 genes with a significant coefficient (at the 0.05 level), 15 had a positive coefficient (all the virulence genes being extra-intestinal VFs) and 2 a negative coefficient (table S21). This suggested that the presence of most of these genes was under positive selection.

To quantify the mode and strength of selection on gene sequences, we computed the ratio of the number of non-synonymous substitutions per non-synonymous site to the number of synonymous substitutions per synonymous site (dN/dS). A ratio above 1 indicates more non-synonymous substitutions than expected under neutral evolution (diversifying selection) and a ration under 1 indicated less non-synonymous substitutions than expected under neutral evolution (purifying selection). We could not compute the dN/dS ratio for *aph*(6)-Id because there was no synonymous mutation. However, the presence of six non-synonymous mutations indicated that this gene was likely under diversifying selection. For virulence genes, we found a significant difference in rates of synonymous and non-synonymous mutations for 17 genes out of 25, with 13 under purifying selection (12 being extra-intestinal VFs, including *kps* operon and *papX*), and 4 under diversifying selection (3 being extra-intestinal VFs, including the *iutA* transmembrane transporter involved in iron uptake) (table S22).

## DISCUSSION

### 1. The phylogenetic history of *E. coli* commensal strains and their serotypes

The primary habitat of *E. coli* is the gut. However, most of the studies interested in virulence and resistance to antibiotics focused on pathogenic collections isolated from extra-intestinal infections, with a few exceptions (35, 32, 36). Here, using whole-genome analysis, we study the phylogenomic evolution of a large commensal collection of *E. coli* (403 strains) over a 30-year period. This collection is unique in its focus on commensals and the timespan, and we controlled as much as we could for covariates.

The proportion of phylogroup B2 strains increased from 9% to 36% while the proportion of phylogroup A strains decreased from 58% to 26% between 1980 and 2010, as already observed in a large part of this collection (14). A predominance of B2 strains over other phylogroups in commensal samples has also been reported in different industrialized countries. For instance, in the late 1990s, the frequency of B2 strains in Australia, Japan, Sweden and USA is between 44 and 48% (1). It is possible that phylogroup B2 increased in frequency just before the 1990s in these industrialized countries as it did in France from 1980 to 2010.

The diversity of STs and serotypes increased over time. The increase in ST diversity was not explained by the increase in frequency of B2 strains (figure S3), but rather by the increase in frequency of rare STs from 2001 to 2010.

We studied in more detail three major STs: two responsible for extra-intestinal infections in humans (ST69 and ST95) (2, 37) and one (ST10) found at high frequency in the human gut as well as in animals (38) (figure 2). The homoplasy (gain or loss of a character independently in different lineages of the tree) of several surface antigens, especially in the ST10 (39), is in agreement with their encoding genes being located in the major hot-spots of recombination in the genome (4). Diversification per time unit of genes encoding for O and H antigens and of FimH protein for ST69 is faster than for ST10 and ST95, which is potentially explained by its younger age as it can be expected during adaptive radiations for example, where lineages diversify rapidly at the emergence of the clade (40).

Several O-groups have been considered as a promising target for a bioconjugate vaccine against extra-intestinal infections (41). Recently, a phase 2 randomized controlled trial showed that a vaccine targeting the O1, O2, O6, and O25-groups was well tolerated and elicited an antibody response against these antigens (8, 9). These four O-groups represented a large part (24%) of our commensal strains. Strains of these four O-groups included STs involved in extra-intestinal infections such as ST69 and ST95, and STs not described as pathogenic clones such as ST10 and ST141. This suggests that the vaccine could perturb the commensal gut microbiota by eliminating commensal *E. coli* clones in addition to the major pathogenic clones.

### 2. Recent temporal evolution of virulence and resistance genes in commensal *E. coli*

Gene and phylogroup frequencies varied in time (figures 1) (14, 34). Strains of phylogroup B2, which carry many VFs, as well as individual VFs increased in frequency, whereas the observed increase in resistance between 1980 and 2001 seems decoupled from phylogroups. To investigate whether virulence and antibiotic resistance gene evolution are governed by different evolutionary mechanisms, we decomposed in two additive components the change in frequency of a gene. A term quantifying the clonal expansion of STs carrying this gene (ST driven change) and a term quantifying the increase in gene frequency within STs, by horizontal transfer following gene duplication (“copy-and-paste” mechanism) or by the increase in frequency of the lineage carrying the focal gene at the expense of other within STs (gene driven change). Their contributions varied between virulence and resistance genes, and between the two periods (1980-2000 and 2000-2010) (figures 4-5).

Between 1980 and 2000, *E. coli* strains rapidly evolved through both the rise of more STs carrying virulence genes and within-ST increase in frequency of virulence genes (gene driven change) (figure 4). In the second time-period from 2001 to 2010, the increase in frequency of STs was mainly driven by the increase in frequency of more virulent STs (ST driven change) (figure 4). Significant changes in frequency suggest a role for direct selection on virulence genes in driving their increase. The presence of several of these genes may be under positive directional selection. Moreover, the sequence diversity of many of them (17 out of 25) also suggests selection, either diversifying or purifying.

What factors could explain the recent increase in virulence genes? Recent environmental changes include a shift towards more processed, and more nutrient-dense food over the period 1969-2002 in France, and a stabilization over 2002-2010 (42), mirroring the spread of virulence genes within ST observed in the period 1980-2000. This could select for virulence genes (e.g., iron acquisition factors) or for virulent STs. Whether bacteria carrying some virulence genes are better adapted to nutrient-dense diets could be experimentally tested (43). In an effort to do so, some of us recently analyzed how mice diet affected the density of *E. coli* in the mice gut and found that the B2 strains used was more prevalent in a high sugar high fat diet than in a high fiber diet (44). In addition to nutriment availability, protection again host immune defense, phages and protozoans may also drive the evolution of virulence genes (2, 45). A change in the diversity and abundance of these predators might likewise explain the recent increase in virulence genes observed in commensal *E. coli*.

For resistance genes, the rapid increase in frequency of several genes [notably *aph*(6)-Id, which is located on plasmids] (46) from 1980 to 2000 was mainly driven by an increase in the frequency of these genes independently within several STs (figure 5). In contrast to 1980-2000, after 2001 the frequency of resistance almost did not change (figure 5). Concomitantly, the proportion of strains resistant to two or more antibiotic categories decreased, indicating a decrease in multi-resistance. In the early 2000s, levels of antibiotic resistance were higher in France than in most European countries (22, 25). A nationwide awareness campaign was launched in 2001 leading to a 26.5% reduction in antibiotic use in humans over 5 years (25). The reduction in antibiotic use could explain the stabilization in resistance gene frequencies observed in our data.

Our work has some limitations. First, only one isolate per subject was sampled. It is well known that populations of subdominant clones are present in the feces (36). Resistant clones are often subdominant and isolated using antibiotic containing plates. They have a specific population structure and epidemiology (47). Second, our data set ends in 2010 and we did not capture the recent evolution. A stool collect campaign is underway and will allow to address this point. Third, the five collections gathered for this study are heterogeneous in terms of host age, sex, location (Paris or Brittany) and potential (but unlikely) antibiotic use. We controlled for these factors in our analyses, and we did not find any large effect of host age, location, sex and the possibility of antibiotic consumption on gene content or genotype. However, we cannot exclude the confounding effects of other unmeasured factors such as diet, travel, comorbidities or contact with farm or pet animals. Fourth, the prevalence of phylogroup B2 and D could have been underestimated because we only considered lactose fermenting isolates (48).

## Conclusion

To investigate the recent evolutionary dynamics of commensal *E. coli*, we whole-genome sequenced a large collection of 403 strains sampled from the stools of healthy volunteers. Several antibiotic resistance genes rapidly spread from 1980 to 2000, largely unhindered by clonal structure. Higher virulence gene frequency evolved through increase in virulence gene frequency within STs and clonal expansion of more virulent STs. Increasing virulence gene frequency of *E. coli* would result, everything else being equal, in an increasing incidence of extra-intestinal infections, and could indeed contribute to the increasing incidence of bacteremia observed in the last decades (12, 15). Future research should investigate whether the observed increasing virulence gene frequency in commensal strains was observed in other geographical areas and what environmental factors could select for *E. coli* virulence genes.

## METHODS

### 1. Strain collections

We studied the whole genomes of four hundred and thirty-six *E. coli* strains gathered from stools of 403 healthy adults (at least 18-year-old) of both sexes living in the Paris area or Brittany (both locations in the North of France) between 1980 to 2010 (see main characteristics of the subjects in table S3). These strains come from five previously published collections: VDG sampled in 1980 (49), ROAR in 2000 (50), LBC in 2001 (51), PAR in 2002 (51), and Coliville in 2010 (table S1) (14). In all studies, one single *E. coli* colony randomly picked on the Drigalski plate was retained per individual (14), probably representing the dominant strain. The study was approved by the ethics evaluation committee of Institut National de la Santé et de la Recherche Médicale (INSERM) (CCTIRS no. 09.243, CNIL no. 909277, and CQI no. 01-014).

In order to improve the temporal span for dating, we used 24 sequences from the Murray collection with samples ranging from 1930 to 1941 (table S23) (52). This collection comprises hundreds of bacterial strains, mostly Enterobacteriaceae, collected between 1917 and 1954 from diverse geographic locations. From the 50 genomic *E. coli* sequences available in the Murray collection, we selected sequences with available sampling time and excluded multiple variants per strain (when strain name, date on tube and origin were identical for two samples).

### 2. Sequencing of the commensal strains

After DNA extraction, whole-genome of each strain was sequenced using Illumina NextSeq 2×150 bp after NextEra XT library preparation (Illumina, San Diego, CA) (26).

### 3. Assembly and typing

The assembly of genomes was performed using the in-house script Petanc that integrates several existing bacterial genomic tools (53), including SPADES (54). This in-house script was also used to perform the typing of strains with several genotyping schemes using the genomic tool SRST2 (55). Multilocus sequence typing (MLST) was performed and STs were defined using the Warwick MLST scheme and the Pasteur scheme (33, 56). We only used the Warwick scheme for the analyses described thereafter. We also determined the O:H types (called serotypes along this manuscript) and the *fimH* alleles (7, 57). The phylogroups were defined using the ClermonTyping method (58). The in-house script Petanc was also used to type the Murray strains in order to determine which strains could be used for dating.

### 4. Distribution of phylogroups, STs and O-groups through time

Because there were three time points close in time with a small number of strains, we aggregated data of years 2000, 2001 and 2002 and it will henceforth be referred to as 2001.

To evaluate the frequency of phylogroups through time, we fitted a binomial generalized linear model (GLM) for the presence/absence of each phylogroup as a function of time (continuous variable), and controlling for several factors heterogeneous over the five collections: host age, location (two categories: Paris vs. Brittany), sex and antibiotic consumption (two categories: “no” vs. “unlikely”). When only an interval was available for age (113 strains from the collections PAR, ROAR and VDG), we used the mean value of this interval. The same analyses were also performed for STs and O-groups.

We generated stacked area charts of phylogroups, STs and O-groups to display their diversity variation between 1980 and 2010.

We analyzed 53 strains in 1980, 104 between 2000 and 2002, and 246 in 2010. To determine whether the number of distinct STs detected each year varied through time or was simply reflecting the number of sampled strains, we generated null distributions of ST number depending on the sampling effort. We sampled 10,000 times a fixed number of strains (53 for 1980, 104 for 2001 and 246 for 2010) with the frequency of each ST set to its overall frequency. For each time point, the observed value was compared to the corresponding null distribution. The same analysis was also performed for serotypes (O and H combinations), O-groups, H-types and *fimH* alleles.

To evaluate the influence of the change in frequency of phylogroup on ST diversity, we generated null distributions of ST number depending on the phylogroup frequency and on the sampling effort. We sampled 10,000 times a fixed number of strains corresponding to the number of strains sampled by phylogroup and by year with the frequency of each ST set to its overall frequency for each phylogroup. For each time point, the observed value was compared to the corresponding null distribution.

### 5. Genomic diversity of the core genome

The 403 assemblies (our commensal collection) were annotated with Prokka (59). We then performed pan-genome analysis from annotated assemblies with Roary on the 403 genomes using default parameters (60). The alignment of the core genome and the list of genes of the accessory genome were generated.

### 6. Core genome phylogeny of E. coli

To build the phylogeny of *E. coli*, we aligned whole genomes (403 strains) to the A phylogroup ST206 R1B5J10 strain with Snippy 4.4.0 using standard parameters (61). We did not remove recombination events because it could accentuate errors in phylogenetic distances and the topology of the tree is usually not affected (62). The alignment of 266,665 SNPs (single-nucleotide polymorphism), was used to produce a maximum likelihood phylogenetic tree with RAxML (63), using the GTRGAMMA model with 1,000 bootstrap replicates.

### 7. Divergence times estimates for major STs

To further study the five most prevalent STs, ST10, ST73, ST95, ST69 and ST59, we aligned whole genomes (136 strains from our collection and 10 from the Murray collection) to references, R1B5J10, 016-002, R1B6J15, H1-004-0023-R-J and IAI39 respectively with Snippy 4.4.0 using standard parameters (61). For each ST, three outgroup sequences were selected as the three closest sequences to the focal ST belonging to two distinct and well supported clades, from the core genome phylogenetic tree. Recombination events were excluded with Gubbins using default settings (64). The five resulting alignments were used to produce a maximum likelihood phylogenetic tree with RAxML using the GTRGAMMA model with 1,000 bootstrap replicates (63).

The temporal analysis was performed with the program BEAST v1.10.4 (65), on three of the most frequent STs with a molecular clock signal (i.e. positive relationship between root-to-tip distance and time): ST10, ST95 and ST69. BEAST implements the “uncorrelated relaxed clock” to model uncorrelated rate variations among lineages; the evolutionary rate of each branch is drawn independently from a common underlying distribution. To estimate divergence times with this model we used the following parameters: uncorrelated log-normal clock, substitution model GTR+I+G, fixed topology (RAxML tree) and 500 million generations with sampling every 1000 generations and a burn-in of 50 million generations. We ran three replicates with three tree priors: coalescent with constant population, coalescent with exponential growth and coalescent with Bayesian skyline. The sample times were used to calibrate the tips. For each dataset, the best fitting model set was determined by computing Bayes factors from marginal likelihoods estimations calculated by stepping-stone sampling (66). Only the models that converged well and had and effective sample size (ESS) larger than 200 for each parameter were compared. The best fitting model was used in the subsequent analyses.

### 8. Virulence and resistance gene repertoire analyses

The resistome, the virulome (both intra- and extra-intestinal virulence genes) and the plasmid type were defined using the in-house script Petanc (53). They were defined by BlastN with Abricate (https://github.com/tseemann/abricate) using the ResFinder database (67), a custom database including the VirulenceFinder database and VFDB (68, 69), to which we added selected genes (table S24), and PlasmidFinder respectively (70). We set the threshold for minimum identity to 80% and for minimum coverage to 90%. If several copies of a gene were detected for a strain, we only kept the copy with the maximal sum of coverage and identity. We also searched for point mutations responsible for betalactam (*ampC* promoter) and fluroquinolone (*gyrA*/*B*, *parC*/*E*) resistance (67). The plasmid sequences were also predicted by PlaScope (71).

To visually explore the relationships between the phylogroups and the resistance and virulence gene repertory we performed a correspondence analysis (CA) (72) with the R package ‘FactoMineR’ (73). A CA is a multivariate graphical analysis used to explore the associations among categorical variables. For each item in a table, a set of factor scores (coordinates) is obtained from linear combinations of rows and columns. These coordinates are projected on two dimensions, with in our case, the first axis opposing the phylogroups with the largest differences. The further the gene categories are from the origin the more they are discriminating. If two variables are close to each other in the plan, they are considered to be strongly associated. The significance of associations between phylogroups and resistance and virulence gene categories was tested with a chi-squared test.

For each isolate, we computed a virulence score corresponding to the number of the virulence factors (VFs) present in each isolate (number of products out of the 104 tested, table S24) (adapted from (74)). The frequency of antibiotic resistance was computed for each year. Both gene acquisitions and point mutations were included (table S25). The corresponding antibiotics were retrieved from ResFinder (75). We next tested whether the virulence score and the frequency of antibiotic resistance changed in time. We fitted a linear model for the number of VF products or for the frequency of antibiotic resistance as a function of year (1980, 2001 (2000-2002), 2010) as a categorical variable.

To decipher whether changes in gene frequency through time were due to variations in ST frequencies or to variations in gene frequencies within STs, we decomposed in two additive components the overall change in gene frequency (*Δ*f*)* as follows (figure S9). We call **f*_i,t_* the frequency of the focal gene in ST *i* at time *t.* We call **p*_i,t_* the frequency of the ST *i* at time *t*. We are interested in the change in frequency of the focal gene from time **t*_1_* to time **t*_2_.* The change in gene frequency can be decomposed in two additive components as such:

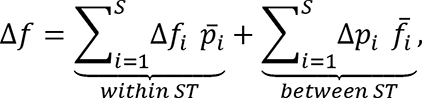

where **S** is the number of STs, *Δ*f*_i_ = *f*_i,t2_ − *f*_i,t1_* is the change in the focal gene frequency in ST *i* from time **t*_1_* to time **t*_2_*, *Δ*p*_i_ = *p*_i,t2_ − *p*_i,t1_* the change in the frequency of ST *i* from time **t*_1_* to time **t*_2_*, 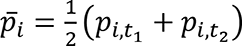 the mean ST *i* frequency in the sample (weighting the two timepoints equally) and 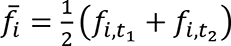 the mean gene frequency in ST *i* in the sample.

For virulence genes, when the frequency of several alleles for a single gene was assessed by our in-house script, we added these frequencies for each corresponding gene (table S24).

The temporal changes of virulence and of antibiotic resistance genes, decomposed in two additive components, were summarized as follow. For virulence genes, we first computed the mean change per product, and then the mean change among product for each category (see table S24). For resistance, we computed the mean change among genes for each category (see table S25).

We next compared the change in frequency of virulence and antibiotic resistance genes to that of other genes in the genome. For each time period, we sampled 1000 non-virulence and non-resistance (NVNR) genes and computed their ST-driven and gene-driven change in frequency as described above. Changes in frequency of virulence and antibiotic resistance genes outside the 95% confidence level computed for NVNR genes were considered significant (0.05 level).

The same analysis (decomposition in two additive components of the change in frequency of virulence and antibiotic resistance genes) was performed for the subset of non-B2 strains only to test if the increase in frequency of virulence genes was due to an overall expansion of the B2 phylogroup.

In addition, we tested if the genes with a significant change in frequency over at least one of the two periods (all virulence or resistance genes with a change in frequency significantly larger than the changes in NVNR genes) were under directional selection over the whole period 1980-2010. More precisely we tested 1) if the presence of these genes (relative to absence) was under selection and 2) if there was a sign of selection acting on the sequence of these genes. First, we fitted a logistic model for gene presence/absence as a function of time (continuous variable, from 1980 to 2020) and phylogroup (categorial variable). Second, we computed dN/dS ratios. We first aligned sequences of each gene with mafft (76). We generated the consensus sequence of each gene (function ‘ConsensusSequence’ in package DECIPHER in R) (77). Then, we calculated dN/dS ratios between each unique sequence and the consensus sequence (function ‘dnds’ in package ape in R) (78). Finally, we tested if the mean dN/dS ratio obtained for each gene was significantly different from 1 (t-test).

## DECLARATIONS

### Ethics approval and consent to participate

Not applicable

### Data availability

The data generated in this study have been submitted to the NCBI BioProject database under the accession numbers PRJEB39252 (Coliville) [https://www.ncbi.nlm.nih.gov/bioproject/?term=PRJEB39252], PRJEB38489 (ROAR) [https://www.ncbi.nlm.nih.gov/bioproject/?term=PRJEB38489], PRJEB44819 (LBC) [https://www.ncbi.nlm.nih.gov/bioproject/?term=PRJEB44819], PRJEB44872 (PAR) [https://www.ncbi.nlm.nih.gov/bioproject/?term=PRJEB44872], PRJEB44873 (VDG) [https://www.ncbi.nlm.nih.gov/bioproject/?term=PRJEB44873].

## Acknowledgements

We thank the reviewers for their valuable suggestions that helped to improve a previous version of this manuscript. We are grateful to Mathieu Thirot for technical assistance. We thank the INRAE MIGALE bioinformatics facility (MIGALE, INRAE, 2020. Migale bioinformatics Facility, doi: 10.15454/1.5572390655343293E12) for providing computing resources. JM and FB were funded by the CNRS Momentum grant to FB. ED was partially supported by the “Fondation pour la Recherche Médicale” (Equipe FRM 2016, grant number DEQ20161136698). GR was supported by a “Poste d’accueil” funded by the Assistance-Publique Hôpitaux de Paris (AP-HP) and the Commissariat à l’Energie Atomique et aux Energies Alternatives (CEA) personal grant for his PhD.

## Appendixes

Tables S1-15 (SupplementalTables_S1_25.xlsx)

Figures S1-11 (SupplementalFigures_S1_11.docx)

## REFERENCES

1. Tenaillon O, Skurnik D, Picard B, Denamur E. 2010. The population genetics of commensal *Escherichia coli*. 3. Nat Rev Microbiol 8:207–217.

2. Denamur E, Clermont O, Bonacorsi S, Gordon D. 2020. The population genetics of pathogenic *Escherichia coli*. 1. Nat Rev Microbiol 19:37–54.

3. Murray CJ, Ikuta KS, Sharara F, Swetschinski L, Aguilar GR, Gray A, Han C, Bisignano C, Rao P, Wool E, Johnson SC, Browne AJ, Chipeta MG, Fell F, Hackett S, Haines-Woodhouse G, Hamadani BHK, Kumaran EAP, McManigal B, Agarwal R, Akech S, Albertson S, Amuasi J, Andrews J, Aravkin A, Ashley E, Bailey F, Baker S, Basnyat B, Bekker A, Bender R, Bethou A, Bielicki J, Boonkasidecha S, Bukosia J, Carvalheiro C, Castañeda-Orjuela C, Chansamouth V, Chaurasia S, Chiurchiù S, Chowdhury F, Cook AJ, Cooper B, Cressey TR, Criollo-Mora E, Cunningham M, Darboe S, Day NPJ, Luca MD, Dokova K, Dramowski A, Dunachie SJ, Eckmanns T, Eibach D, Emami A, Feasey N, Fisher-Pearson N, Forrest K, Garrett D, Gastmeier P, Giref AZ, Greer RC, Gupta V, Haller S, Haselbeck A, Hay SI, Holm M, Hopkins S, Iregbu KC, Jacobs J, Jarovsky D, Javanmardi F, Khorana M, Kissoon N, Kobeissi E, Kostyanev T, Krapp F, Krumkamp R, Kumar A, Kyu HH, Lim C, Limmathurotsakul D, Loftus MJ, Lunn M, Ma J, Mturi N, Munera-Huertas T, Musicha P, Mussi-Pinhata MM, Nakamura T, Nanavati R, Nangia S, Newton P, Ngoun C, Novotney A, Nwakanma D, Obiero CW, Olivas-Martinez A, Olliaro P, Ooko E, Ortiz-Brizuela E, Peleg AY, Perrone C, Plakkal N, Ponce-de-Leon A, Raad M, Ramdin T, Riddell A, Roberts T, Robotham JV, Roca A, Rudd KE, Russell N, Schnall J, Scott JAG, Shivamallappa M, Sifuentes-Osornio J, Steenkeste N, Stewardson AJ, Stoeva T, Tasak N, Thaiprakong A, Thwaites G, Turner C, Turner P, Doorn HR van, Velaphi S, Vongpradith A, Vu H, Walsh T, Waner S, Wangrangsimakul T, Wozniak T, Zheng P, Sartorius B, Lopez AD, Stergachis A, Moore C, Dolecek C, Naghavi M. 2022. Global burden of bacterial antimicrobial resistance in 2019: a systematic analysis. The Lancet 399:629–655.

4. Touchon M, Hoede C, Tenaillon O, Barbe V, Baeriswyl S, Bidet P, Bingen E, Bonacorsi S, Bouchier C, Bouvet O. 2009. Organised genome dynamics in the *Escherichia coli* species results in highly diverse adaptive paths. PLoS Genet 5.

5. Orskov I, Orskov F, Jann B, Jann K. 1977. Serology, chemistry, and genetics of O and K antigens of *Escherichia coli*. Bacteriol Rev 41:667–710.

6. Schembri MA, Kjaergaard K, Sokurenko EV, Klemm P. 2001. Molecular characterization of the *Escherichia coli* FimH adhesin. J Infect Dis 183:S28–S31.

7. Roer L, Tchesnokova V, Allesøe R, Muradova M, Chattopadhyay S, Ahrenfeldt J, Thomsen MCF, Lund O, Hansen F, Hammerum AM, Sokurenko E, Hasman H. 2017. Development of a web tool for *Escherichia coli* subtyping based on fimH alleles. J Clin Microbiol 55:2538–2543.

8. Frenck RW, Ervin J, Chu L, Abbanat D, Spiessens B, Go O, Haazen W, van den Dobbelsteen G, Poolman J, Thoelen S, Ibarra de Palacios P. 2019. Safety and immunogenicity of a vaccine for extra-intestinal pathogenic *Escherichia coli* (ESTELLA): a phase 2 randomised controlled trial. Lancet Infect Dis 19:631–640.

9. Huttner A, Hatz C, van den Dobbelsteen G, Abbanat D, Hornacek A, Frölich R, Dreyer AM, Martin P, Davies T, Fae K, van den Nieuwenhof I, Thoelen S, de Vallière S, Kuhn A, Bernasconi E, Viereck V, Kavvadias T, Kling K, Ryu G, Hülder T, Gröger S, Scheiner D, Alaimo C, Harbarth S, Poolman J, Fonck VG. 2017. Safety, immunogenicity, and preliminary clinical efficacy of a vaccine against extraintestinal pathogenic *Escherichia coli* in women with a history of recurrent urinary tract infection: a randomised, single-blind, placebo-controlled phase 1b trial. Lancet Infect Dis 17:528–537.

10. Johnson JR. 1991. Virulence factors in *Escherichia coli* urinary tract infection. Clin Microbiol Rev 4:80–128.

11. Clermont O, Couffignal C, Blanco J, Mentré F, Picard B, Denamur E, Groups the C and C. 2017. Two levels of specialization in bacteraemic *Escherichia coli* strains revealed by their comparison with commensal strains. Epidemiol Infect 145:872–882.

12. Kraker MEA de, Davey PG, Grundmann H, Group on behalf of the B study. 2011. Mortality and hospital stay associated with resistant *Staphylococcus aureus* and *Escherichia coli* bacteremia: estimating the burden of antibiotic resistance in Europe. PLOS Med 8:e1001104.

13. Didelot X, Darling A, Falush D. 2009. Inferring genomic flux in bacteria. Genome Res 19:306–317.

14. Massot M, Daubié A-S, Clermont O, Jaureguy F, Couffignal C, Dahbi G, Mora A, Blanco J, Branger C, Mentré F, Eddi A, Picard B, Denamur E. 2016. Phylogenetic, virulence and antibiotic resistance characteristics of commensal strain populations of *Escherichia coli* from community subjects in the Paris area in 2010 and evolution over 30 years. Microbiology 162:642–650.

15. Vihta K-D, Stoesser N, Llewelyn MJ, Quan TP, Davies T, Fawcett NJ, Dunn L, Jeffery K, Butler CC, Hayward G, Andersson M, Morgan M, Oakley S, Mason A, Hopkins S, Wyllie DH, Crook DW, Wilcox MH, Johnson AP, Peto TEA, Walker AS. 2018. Trends over time in *Escherichia coli* bloodstream infections, urinary tract infections, and antibiotic susceptibilities in Oxfordshire, UK, 1998–2016: a study of electronic health records. Lancet Infect Dis 18:1138–1149.

16. Birgy A, Levy C, Bidet P, Thollot F, Derkx V, Béchet S, Mariani-Kurkdjian P, Cohen R, Bonacorsi S. 2016. ESBL-producing Escherichia coli ST131 versus non-ST131: evolution and risk factors of carriage among French children in the community between 2010 and 2015. J Antimicrob Chemother 71:2949–2956.

17. Le Gall T, Clermont O, Gouriou S, Picard B, Nassif X, Denamur E, Tenaillon O. 2007. Extraintestinal virulence is a coincidental by-product of commensalism in B2 phylogenetic group *Escherichia coli* strains. Mol Biol Evol 24:2373–2384.

18. Diard M, Garry L, Selva M, Mosser T, Denamur E, Matic I. 2010. Pathogenicity-associated islands in extraintestinal pathogenic *Escherichia coli* are fitness elements involved in intestinal colonization. J Bacteriol 192:4885–4893.

19. Östblom A, Adlerberth I, Wold AE, Nowrouzian FL. 2011. Pathogenicity island markers, virulence determinants malX and usp, and the capacity of *Escherichia coli* to persist in infants’ commensal microbiotas. Appl Environ Microbiol 77:2303–2308.

20. Nicolas-Chanoine M-H, Bertrand X, Madec J-Y. 2014. *Escherichia coli* ST131, an intriguing clonal group. Clin Microbiol Rev 27:543–574.

21. Martin P, Marcq I, Magistro G, Penary M, Garcie C, Payros D, Boury M, Olier M, Nougayrède J-P, Audebert M, Chalut C, Schubert S, Oswald E. 2013. Interplay between siderophores and colibactin genotoxin biosynthetic pathways in *Escherichia coli*. PLOS Pathog 9:e1003437.

22. Goossens H, Ferech M, Vander Stichele R, Elseviers M. 2005. Outpatient antibiotic use in Europe and association with resistance: a cross-national database study. The Lancet 365:579–587.

23. Chatterjee A, Modarai M, Naylor NR, Boyd SE, Atun R, Barlow J, Holmes AH, Johnson A, Robotham JV. 2018. Quantifying drivers of antibiotic resistance in humans: a systematic review. Lancet Infect Dis 18:e368–e378.

24. Tedijanto C, Olesen SW, Grad YH, Lipsitch M. 2018. Estimating the proportion of bystander selection for antibiotic resistance among potentially pathogenic bacterial flora. Proc Natl Acad Sci 115:E11988–E11995.

25. Sabuncu E, David J, Bernède-Bauduin C, Pépin S, Leroy M, Boëlle P-Y, Watier L, Guillemot D. 2009. Significant reduction of antibiotic use in the community after a nationwide campaign in France, 2002–2007. PLoS Med 6:e1000084.

26. de Lastours V, Laouénan C, Royer G, Carbonnelle E, Lepeule R, Esposito-Farèse M, Clermont O, Duval X, Fantin B, Mentré F, Decousser JW, Denamur E, Lefort A, Group on behalf of the S. 2020. Mortality in *Escherichia coli* bloodstream infections: antibiotic resistance still does not make it. J Antimicrob Chemother 75:2334–2343.

27. Kallonen T, Brodrick HJ, Harris SR, Corander J, Brown NM, Martin V, Peacock SJ, Parkhill J. 2017. Systematic longitudinal survey of invasive *Escherichia coli* in England demonstrates a stable population structure only transiently disturbed by the emergence of ST131. Genome Res 27:1437–1449.

28. Salipante SJ, Roach DJ, Kitzman JO, Snyder MW, Stackhouse B, Butler-Wu SM, Lee C, Cookson BT, Shendure J. 2015. Large-scale genomic sequencing of extraintestinal pathogenic *Escherichia coli* strains. Genome Res 25:119–128.

29. Foxman B. 2010. The epidemiology of urinary tract infection. 12. Nat Rev Urol 7:653–660.

30. Goto M, McDanel JS, Jones MM, Livorsi DJ, Ohl ME, Beck BF, Richardson KK, Alexander B, Perencevich EN. 2017. Antimicrobial nonsusceptibility of gram-negative bloodstream isolates, Veterans Health Administration System, United States, 2003–20131. Emerg Infect Dis 23:1815–1825.

31. Kauffmann F. 1947. The serology of the *coli* group. J Immunol Baltim Md 1950 57:71–100.

32. Arimizu Y, Kirino Y, Sato MP, Uno K, Sato T, Gotoh Y, Auvray F, Brugere H, Oswald E, Mainil JG, Anklam KS, Döpfer D, Yoshino S, Ooka T, Tanizawa Y, Nakamura Y, Iguchi A, Morita-Ishihara T, Ohnishi M, Akashi K, Hayashi T, Ogura Y. 2019. Large-scale genome analysis of bovine commensal *Escherichia coli* reveals that bovine-adapted *E. coli* lineages are serving as evolutionary sources of the emergence of human intestinal pathogenic strains. Genome Res 29:1495–1505.

33. Wirth T, Falush D, Lan R, Colles F, Mensa P, Wieler LH, Karch H, Reeves PR, Maiden MCJ, Ochman H, Achtman M. 2006. Sex and virulence in *Escherichia coli*: an evolutionary perspective. Mol Microbiol 60:1136–1151.

34. Touchon M, Perrin A, de Sousa JAM, Vangchhia B, Burn S, O’Brien CL, Denamur E, Gordon D, Rocha EP. 2020. Phylogenetic background and habitat drive the genetic diversification of *Escherichia coli*. PLOS Genet 16:e1008866.

35. Qin X, Hu F, Wu S, Ye X, Zhu D, Zhang Y, Wang M. 2013. Comparison of adhesin genes and antimicrobial susceptibilities between uropathogenic and intestinal commensal *Escherichia coli* strains. PLOS ONE 8:e61169.

36. Smati M, Clermont O, Gal FL, Schichmanoff O, Jauréguy F, Eddi A, Denamur E, Picard B, Group for the C. 2013. Real-time PCR for quantitative analysis of human commensal *Escherichia coli* populations reveals a high frequency of subdominant phylogroups. Appl Environ Microbiol 79:5005–5012.

37. Basmaci R, Bonacorsi S, Bidet P, Biran V, Aujard Y, Bingen E, Béchet S, Cohen R, Levy C. 2015. *Escherichia coli* meningitis features in 325 children from 2001 to 2013 in France. Clin Infect Dis 61:779–786.

38. Manges AR, Harel J, Masson L, Edens TJ, Portt A, Reid-Smith RJ, Zhanel GG, Kropinski AM, Boerlin P. 2015. Multilocus sequence typing and virulence gene profiles associated with *Escherichia coli* from human and animal sources. Foodborne Pathog Dis 12:302–310.

39. Royer G, Darty MM, Clermont O, Condamine B, Laouenan C, Decousser J-W, Vallenet D, Lefort A, de Lastours V, Denamur E, Wolff M, Alavoine L, Duval X, Skurnik D, Woerther P-L, Andremont A, Carbonnelle E, Lortholary O, Nassif X, Abgrall S, Jaureguy F, Picard B, Houdouin V, Aujard Y, Bonacorsi S, Meybeck A, Barnaud G, Branger C, Lefort A, Fantin B, Bellier C, Bert F, Nicolas-Chanoine M-H, Page B, Cremniter J, Gaillard J-L, Leturdu F, Sollet J-P, Plantefève G, Panhard X, Mentré F, Marcault E, Tubach F, Zarrouk V, Bert F, Duprilot M, Leflon-Guibout V, Maataoui N, Armand L, Nguyen LL, Collarino R, Munier A-L, Jacquier H, Lecorché E, Coutte L, Gomart C, Fateh OA, Landraud L, Messika J, Aslangul E, Gerin M, Bleibtreu A, Lescat M, Walewski V, Mechaï F, Dollat M, Maherault A-C, Wolff M, Mercier-Darty M, Basse B, COLIBAFI and SEPTICOLI groups. 2021. Phylogroup stability contrasts with high within sequence type complex dynamics of *Escherichia coli* bloodstream infection isolates over a 12-year period. Genome Med 13:77.

40. Barrier M, Robichaux RH, Purugganan MD. 2001. Accelerated regulatory gene evolution in an adaptive radiation. Proc Natl Acad Sci 98:10208–10213.

41. Poolman JT, Wacker M. 2016. Extraintestinal pathogenic *Escherichia coli*, a common human pathogen: challenges for vaccine development and progress in the field. J Infect Dis 213:6–13.

42. Caillavet F, Darmon N, Létoile F, Nichèle V. 2018. Is nutritional quality of food-at-home purchases improving? 1969–2010: 40 years of household consumption surveys in France. Eur J Clin Nutr 72:220–227.

43. O’Brien CL, Gordon DMY 2011. 2011. Effect of diet and gut dynamics on the establishment and persistence of Escherichia coli. Microbiology 157:1375–1384.

44. Ghalayini M, Magnan M, Dion S, Zatout O, Bourguignon L, Tenaillon O, Lescat M. 2019. Long-term evolution of the natural isolate of *Escherichia coli* 536 in the mouse gut colonized after maternal transmission reveals convergence in the constitutive expression of the lactose operon. Mol Ecol 28:4470–4485.

45. Wildschutte H, Wolfe DM, Tamewitz A, Lawrence JG. 2004. Protozoan predation, diversifying selection, and the evolution of antigenic diversity in *Salmonella*. Proc Natl Acad Sci 101:10644–10649.

46. Khezri A, Avershina E, Ahmad R. 2021. Plasmid identification and plasmid-mediated antimicrobial gene detection in Norwegian isolates. 1. Microorganisms 9:52.

47. Day MJ, Hopkins KL, Wareham DW, Toleman MA, Elviss N, Randall L, Teale C, Cleary P, Wiuff C, Doumith M, Ellington MJ, Woodford N, Livermore DM. 2019. Extended-spectrum β-lactamase-producing *Escherichia coli* in human-derived and foodchain-derived samples from England, Wales, and Scotland: an epidemiological surveillance and typing study. Lancet Infect Dis 19:1325–1335.

48. Chakraborty A, Adhikari P, Shenoy S, Saralaya V. 2016. Virulence factor profiles, phylogenetic background, and antimicrobial resistance pattern of lactose fermenting and nonlactose fermenting *Escherichia coli* from extraintestinal sources. Indian J Pathol Microbiol 59:180–184

49. Duriez P, Clermont O, Bonacorsi S, Bingen E, Chaventré A, Elion J, Picard B, Denamur E. 2001. Commensal *Escherichia coli* isolates are phylogenetically distributed among geographically distinct human populations. Microbiology, 147:1671–1676.

50. Skurnik D, Clermont O, Guillard T, Launay A, Danilchanka O, Pons S, Diancourt L, Lebreton F, Kadlec K, Roux D, Jiang D, Dion S, Aschard H, Denamur M, Cywes-Bentley C, Schwarz S, Tenaillon O, Andremont A, Picard B, Mekalanos J, Brisse S, Denamur E. 2016. Emergence of antimicrobial-resistant *Escherichia coli* of animal origin spreading in humans. Mol Biol Evol 33:898–914.

51. Escobar-Páramo P, Grenet K, Menac’h AL, Rode L, Salgado E, Amorin C, Gouriou S, Picard B, Rahimy MC, Andremont A, Denamur E, Ruimy R. 2004. Large-scale population Structure of human commensal *Escherichia coli* isolates. Appl Environ Microbiol 70:5698–5700.

52. Baker KS, Burnett E, McGregor H, Deheer-Graham A, Boinett C, Langridge GC, Wailan AM, Cain AK, Thomson NR, Russell JE, Parkhill J. 2015. The Murray collection of pre-antibiotic era Enterobacteriacae: a unique research resource. Genome Med 7:97.

53. Bourrel AS, Poirel L, Royer G, Darty M, Vuillemin X, Kieffer N, Clermont O, Denamur E, Nordmann P, Decousser J-W, IAME Resistance Group. 2019. Colistin resistance in Parisian inpatient faecal *Escherichia coli* as the result of two distinct evolutionary pathways. J Antimicrob Chemother 74:1521–1530.

54. Bankevich A, Nurk S, Antipov D, Gurevich AA, Dvorkin M, Kulikov AS, Lesin VM, Nikolenko SI, Pham S, Prjibelski AD, Pyshkin AV, Sirotkin AV, Vyahhi N, Tesler G, Alekseyev MA, Pevzner PA. 2012. SPAdes: A new genome assembly algorithm and its applications to single-cell sequencing. J Comput Biol 19:455–477.

55. Inouye M, Dashnow H, Raven L-A, Schultz MB, Pope BJ, Tomita T, Zobel J, Holt KE. 2014. SRST2: Rapid genomic surveillance for public health and hospital microbiology labs. Genome Med 6:90.

56. Jaureguy F, Landraud L, Passet V, Diancourt L, Frapy E, Guigon G, Carbonnelle E, Lortholary O, Clermont O, Denamur E, Picard B, Nassif X, Brisse S. 2008. Phylogenetic and genomic diversity of human bacteremic *Escherichia coli* strains. BMC Genomics 9:560.

57. Ingle DJ, Valcanis M, Kuzevski A, Tauschek M, Inouye M, Stinear T, Levine MM, Robins-Browne RM, Holt KE. 2016. In silico serotyping of E. coli from short read data identifies limited novel O-loci but extensive diversity of O:H serotype combinations within and between pathogenic lineages. Microb Genomics 2.

58. Beghain J, Bridier-Nahmias A, Le Nagard H, Denamur E, Clermont O. 2018. ClermonTyping: an easy-to-use and accurate in silico method for *Escherichia genus* strain phylotyping. Microb Genomics 4.

59. Seemann T. 2014. Prokka: rapid prokaryotic genome annotation. Bioinformatics 30:2068–2069.

60. Page AJ, Cummins CA, Hunt M, Wong VK, Reuter S, Holden MTG, Fookes M, Falush D, Keane JA, Parkhill J. 2015. Roary: rapid large-scale prokaryote pan genome analysis. Bioinformatics 31:3691–3693.

61. Seemann T. 2015. Snippy: rapid haploid variant calling and core SNP phylogeny. GitHub Available Github Comtseemannsnippy.

62. Hedge J, Wilson DJ. 2014. Bacterial phylogenetic reconstruction from whole genomes is robust to recombination but demographic inference is not. mBio 5.

63. Stamatakis A. 2014. RAxML version 8: a tool for phylogenetic analysis and post-analysis of large phylogenies. Bioinformatics 30:1312–1313.

64. Croucher NJ, Page AJ, Connor TR, Delaney AJ, Keane JA, Bentley SD, Parkhill J, Harris SR. 2015. Rapid phylogenetic analysis of large samples of recombinant bacterial whole genome sequences using Gubbins. Nucleic Acids Res 43:e15–e15.

65. Drummond AJ, Suchard MA, Xie D, Rambaut A. 2012. Bayesian phylogenetics with BEAUti and the BEAST 1.7. Mol Biol Evol 29:1969–1973.

66. Baele G, Lemey P, Suchard MA. 2016. Genealogical working distributions for bayesian model testing with phylogenetic uncertainty. Syst Biol 65:250–264.

67. Zankari E, Hasman H, Cosentino S, Vestergaard M, Rasmussen S, Lund O, Aarestrup FM, Larsen MV. 2012. Identification of acquired antimicrobial resistance genes. J Antimicrob Chemother 67:2640–2644.

68. Chen L, Zheng D, Liu B, Yang J, Jin Q. 2016. VFDB 2016: hierarchical and refined dataset for big data analysis--10 years on. Nucleic Acids Res 44:D694–697.

69. Joensen KG, Scheutz F, Lund O, Hasman H, Kaas RS, Nielsen EM, Aarestrup FM. 2014. Real-time whole-genome sequencing for routine typing, surveillance, and outbreak detection of verotoxigenic *Escherichia coli*. J Clin Microbiol 52:1501–1510.

70. Carattoli A, Zankari E, García-Fernández A, Voldby Larsen M, Lund O, Villa L, Møller Aarestrup F, Hasman H. 2014. *In Silico* detection and typing of plasmids using PlasmidFinder and Plasmid multilocus sequence typing. Antimicrob Agents Chemother 58:3895–3903.

71. Royer G, Decousser JW, Branger C, Médigue C, Denamur E, Vallenet D. 2018. PlaScope: a targeted approach to assess the plasmidome of *Escherichia coli* strains. preprint, Bioinformatics.

72. Benzecri J. 1992. Correspondence analysis handbook. Marcel Decker, New York.

73. Lê S, Josse J, Husson F. 2008. FactoMineR : An *R* Package for multivariate analysis. J Stat Softw 25.

74. Lefort A, Panhard X, Clermont O, Woerther P-L, Branger C, Mentré F, Fantin B, Wolff M, Denamur E. 2011. Host factors and portal of entry outweigh bacterial determinants to predict the severity of *Escherichia coli* bacteremia. J Clin Microbiol 49:777–783.

75. Bortolaia V, Kaas RS, Ruppe E, Roberts MC, Schwarz S, Cattoir V, Philippon A, Allesoe RL, Rebelo AR, Florensa AF, Fagelhauer L, Chakraborty T, Neumann B, Werner G, Bender JK, Stingl K, Nguyen M, Coppens J, Xavier BB, Malhotra-Kumar S, Westh H, Pinholt M, Anjum MF, Duggett NA, Kempf I, Nykäsenoja S, Olkkola S, Wieczorek K, Amaro A, Clemente L, Mossong J, Losch S, Ragimbeau C, Lund O, Aarestrup FM. 2020. ResFinder 4.0 for predictions of phenotypes from genotypes. J Antimicrob Chemother 75:3491–3500.

76. Katoh K, Standley DM. 2013. MAFFT Multiple sequence alignment software Version 7: Improvements in performance and usability. Mol Biol Evol 30:772–780.

77. Wright E S. 2016. Using DECIPHER v2.0 to Analyze big biological sequence data in R. R J 8:352.

78. Paradis E, Schliep K. 2019. ape 5.0: an environment for modern phylogenetics and evolutionary analyses in R. Bioinformatics 35:526–528.

