## Supplemental Figures for "The population genomics of increased virulence and antibiotic resistance in human commensal *Escherichia coli* over 30 years in France"

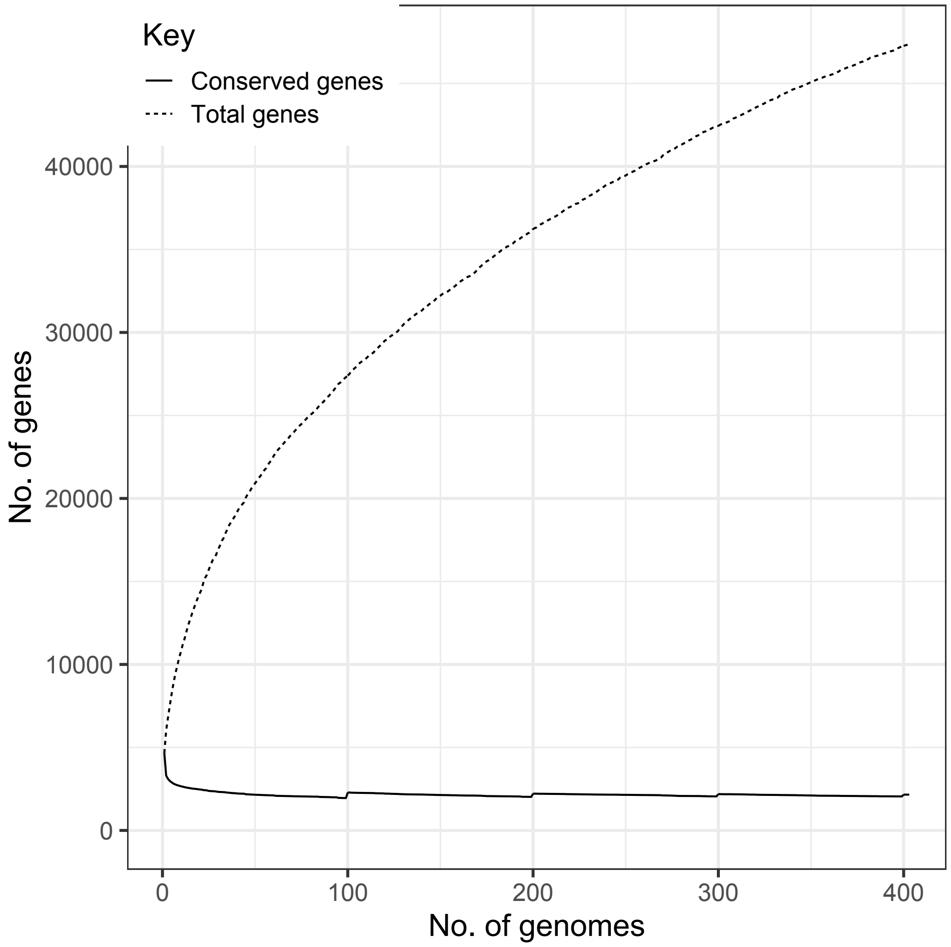


**Figure S1.** Pan genome variation with the number of genomes analysed (Roary output). No.: number.

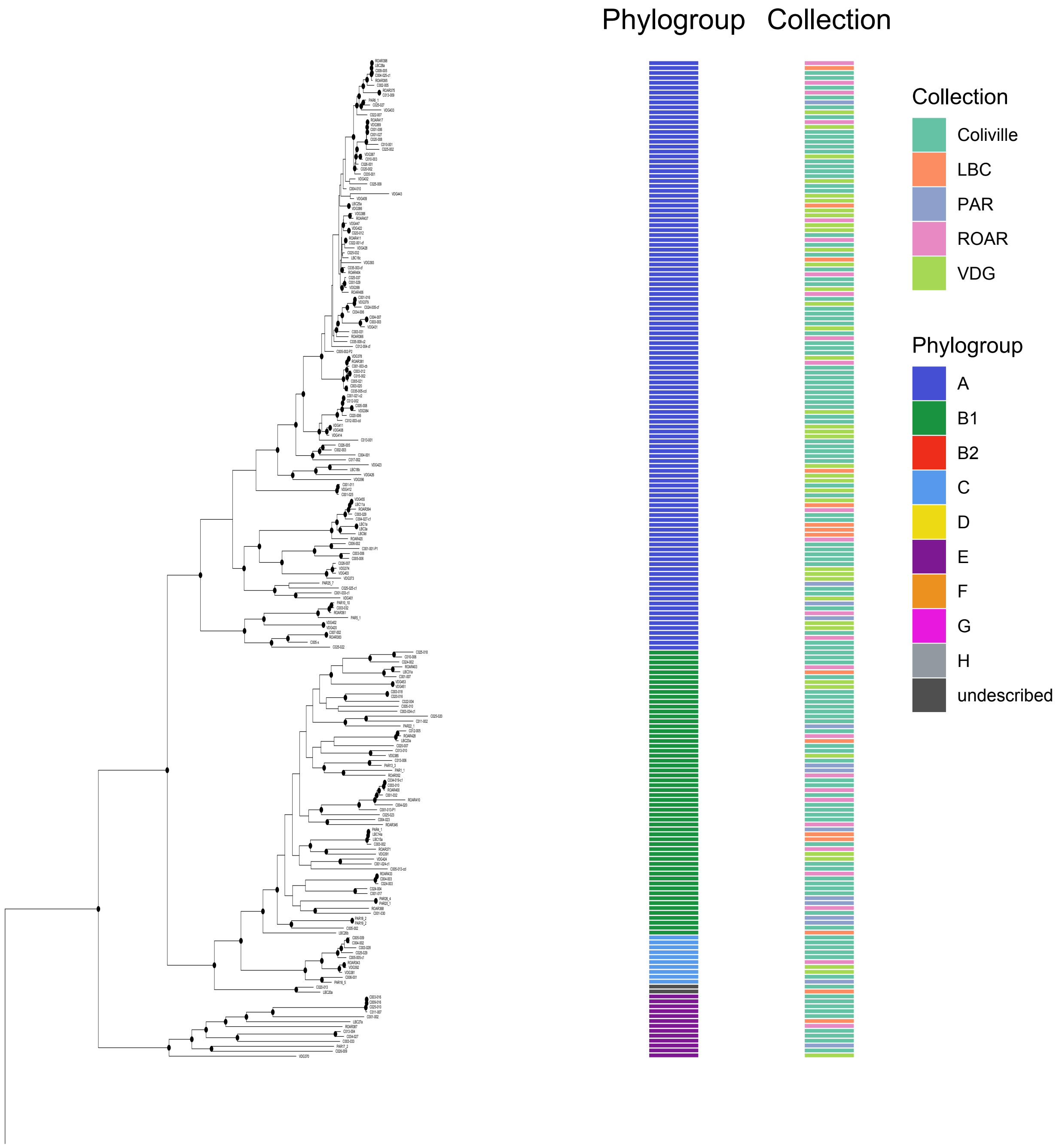


**Figure S2 (part 1)** *E. coli* phylogenetic tree built with RAxML. The tree is rooted on the isolates belonging to the clades IV and V (3 isolates) that are not shown here. Nodes with a bootstrap support > 90 are indicated with black circles.

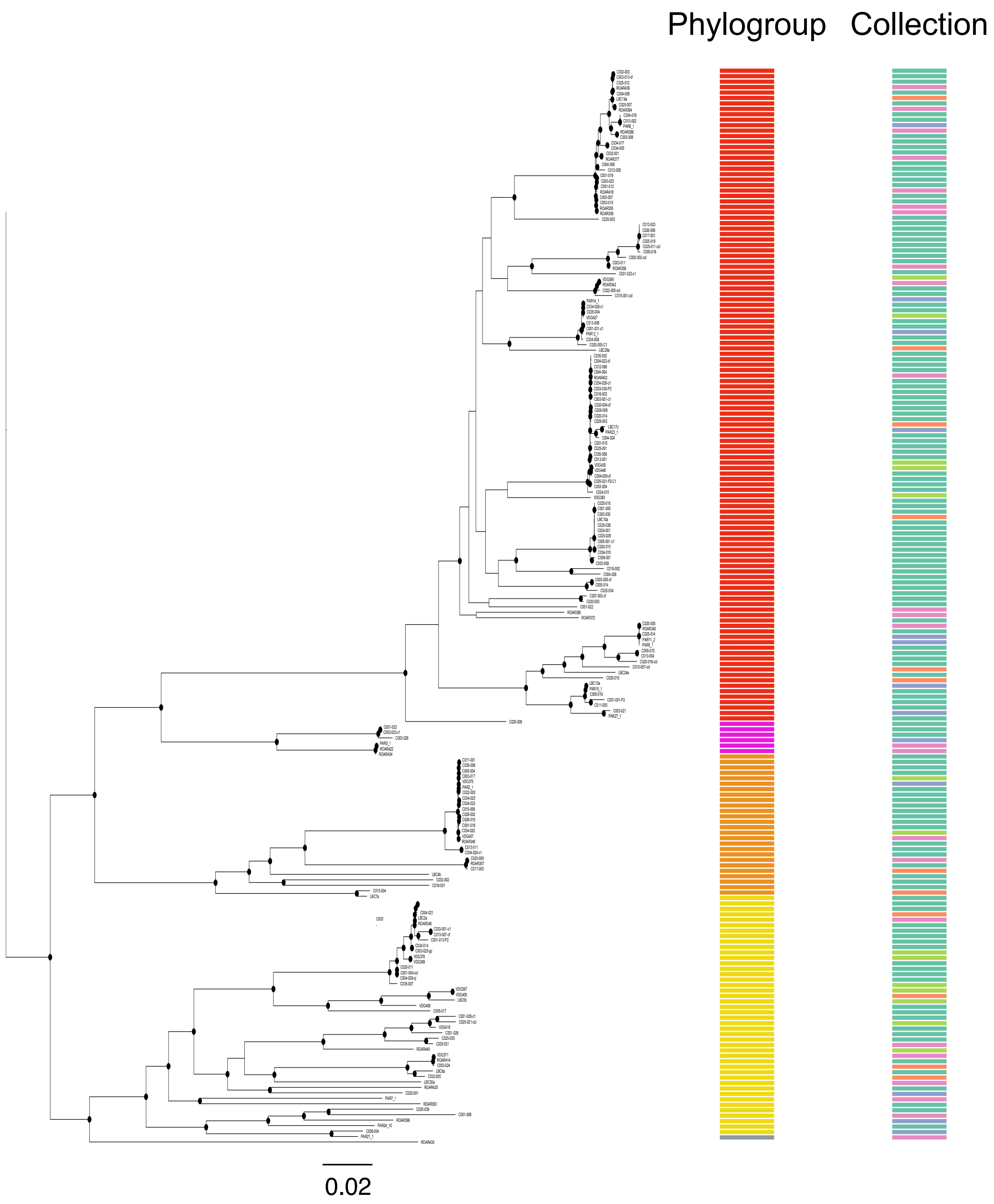


**Figure S2 (part 2)** *E. coli* phylogenetic tree built with RAxML. The tree is rooted on the isolates belonging to the clades IV and V (3 isolates) that are not shown here. Nodes with a bootstrap support > 90 are indicated with black circles.

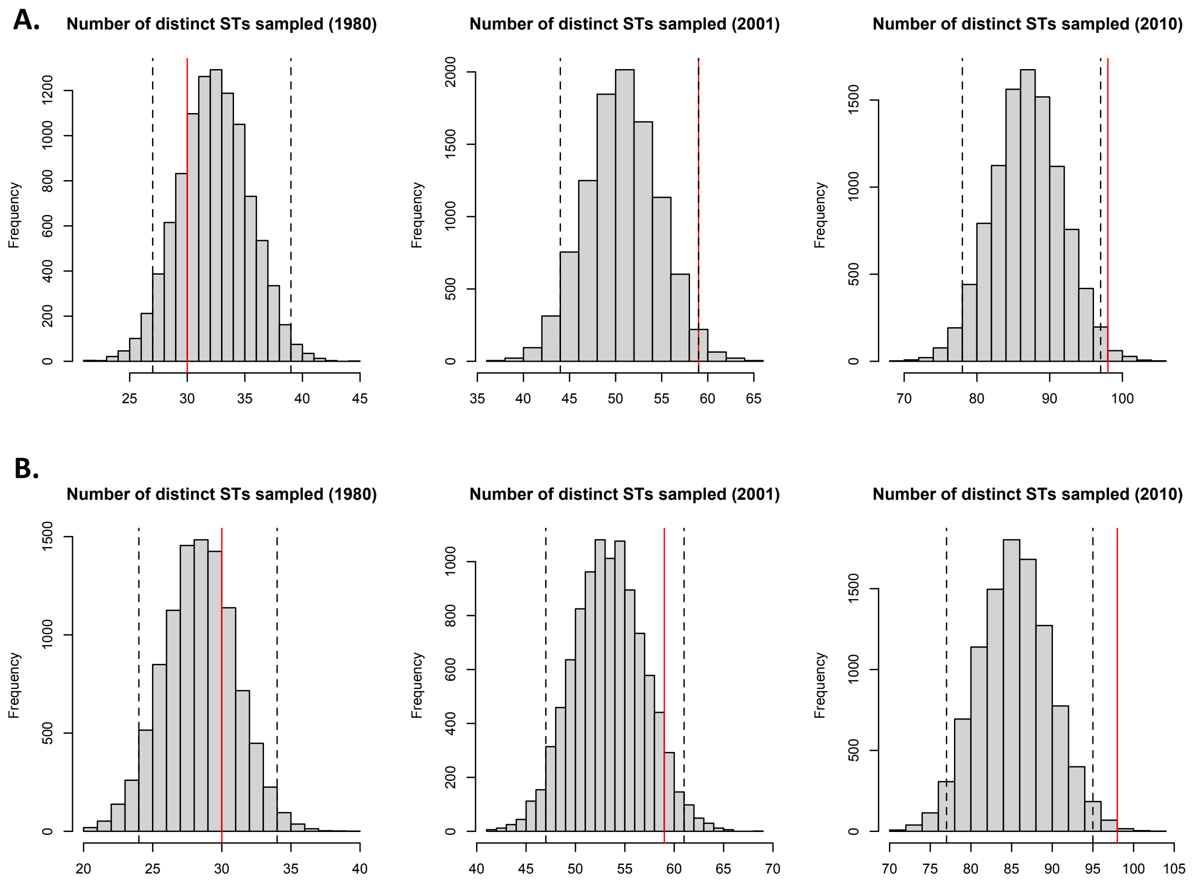


**Figure S3.** (A) Null distributions of the number of distinct STs sampled depending on the strain sample size. STs were sampled depending on the overall frequency of each ST in the collection. (B) Null distributions of the number of distinct STs sampled depending on the phylogroup frequencies and on the strain sample size. STs were sampled depending on the overall frequency of each ST in each phylogroup. The red line is the observed number of STs for each year (1980, 2001 (2000-2002) and 2010). The dashed lines represent the 95% prediction interval.

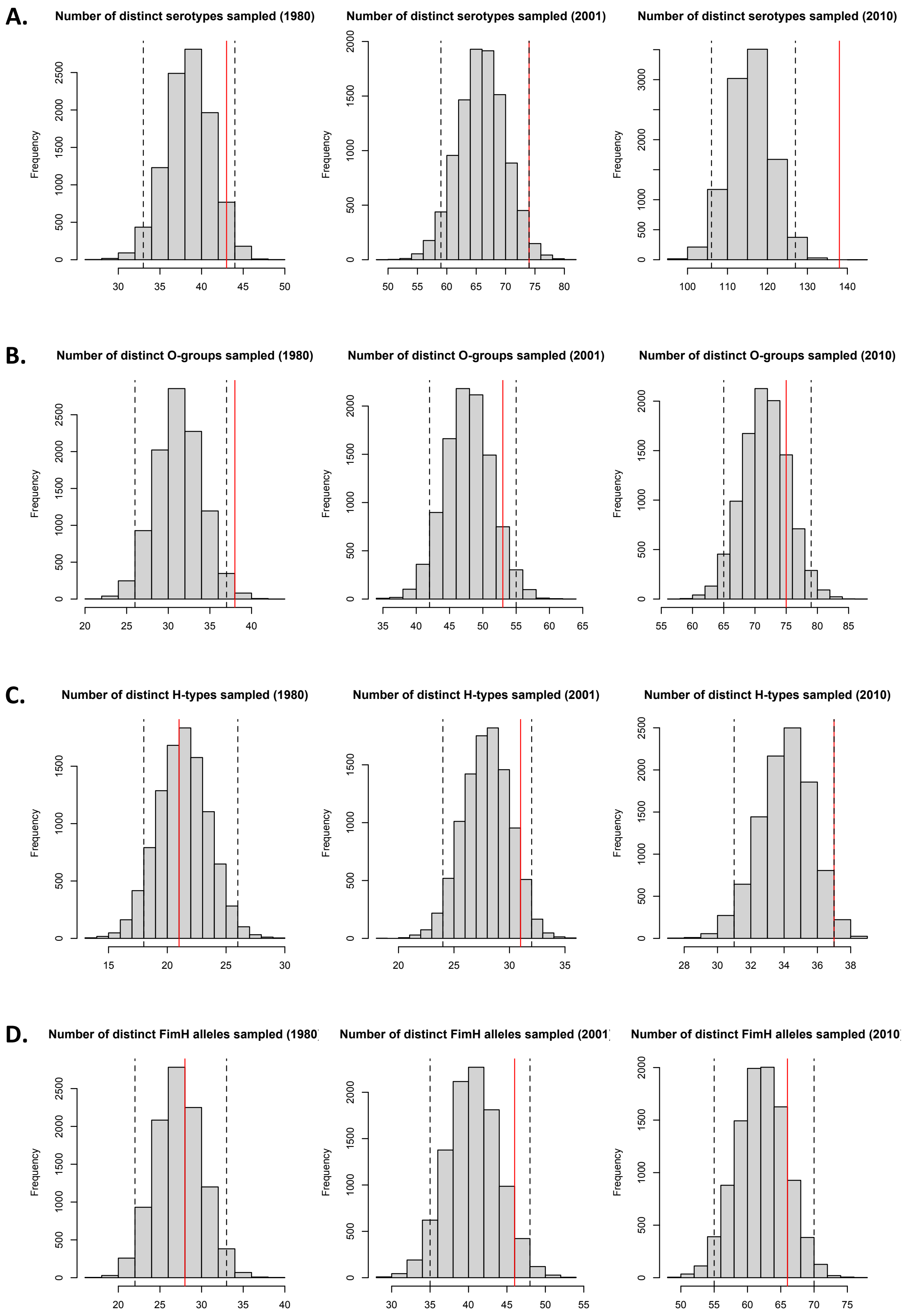


**Figure S4.** Null distributions of the number of distinct (A) serotypes (O and H combinations), (B) O-groups, (C) H-types and (D) FimH alleles sampled depending on the strain sample size. Serotypes, O-groups, H-types and FimH alleles were sampled depending on the overall frequency of each serotype, O-group, H-type and FimH allele respectively in the collection. The red line is the observed number of serotypes, O-groups, H-types and FimH alleles for each year (1980, 2001 (2000-2002) and 2010). The dashed lines represent the 95% prediction interval.

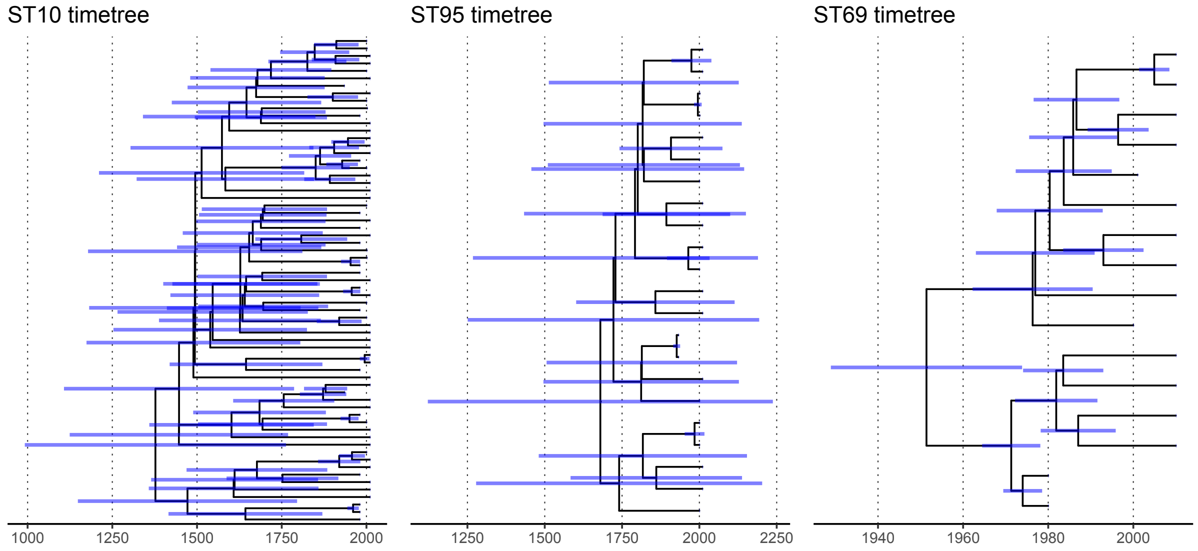


**Figure S5.** Timetrees of ST10, ST95 and ST69 with confidence intervals. Divergence times were estimated with BEAST v1.10.4 (Drummond et al. 2012).

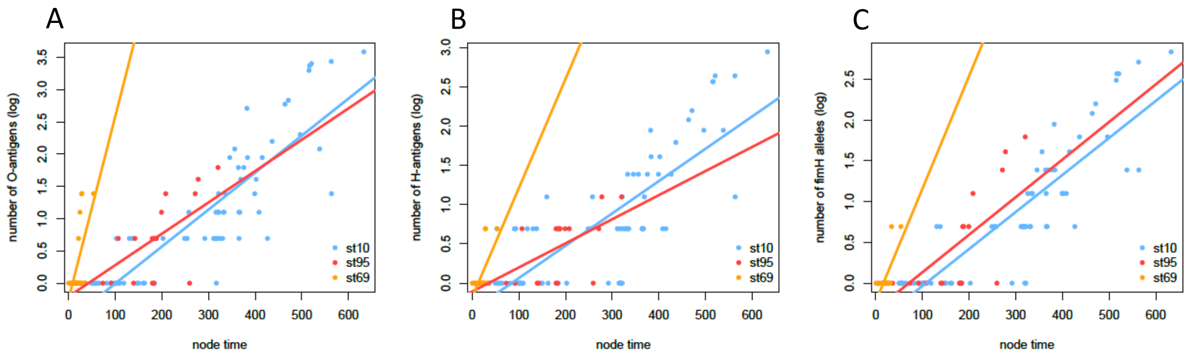


**Figure S6**. Relationship between O-antigen (A), H-antigen (B) and *fimH* allele (C) diversity and node age (years) for ST10, ST95 and ST69. We computed the number of distinct antigens and alleles associated to each node of the trees and fitted a linear model for the node log-transformed number of antigens (O or H) or *fimH* alleles as a function of node age, for each ST tree. We found a positive relationship between the log-transformed number of serotypes and node age with a similar coefficient for ST10 (O-antigen: slope=5.73e-03, p-value<2.2e-16; H-antigen: slope=4.11e-03, p-value=8.57e-16; *fimH*: slope=4.53e-03, p-value<2.2e-16) and ST95 (O-antigen: slope=4.86e-03, p-value=8.20e-05; H-antigen: slope=3.07e-03, p-value=3.40e-04; *fimH*: slope=4.60e-03, p-value=1.83e-04) and a higher coefficient for ST69 (O-antigen: slope=2.83e-02, p-value=2.54e-02; H-antigen: slope=1.39e-02, p-value=2.01e-02; *fimH*: slope=1.38e-02, p-value=4.21e-03).

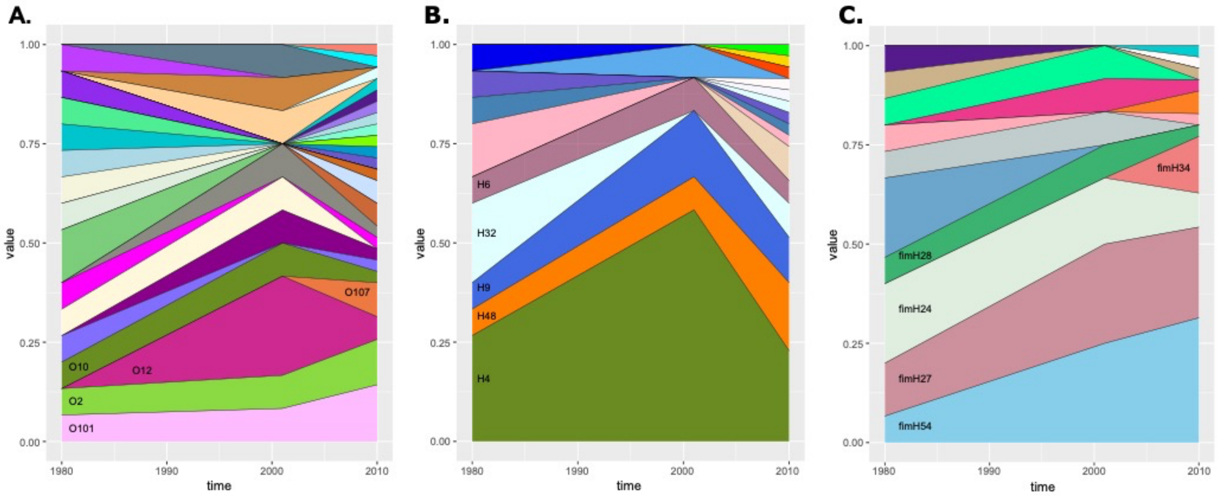


**Figure S7.** Proportion of O-antigens (A), H-antigens (B) and *fimH* alleles (C) between 1980 and 2010 among the 62 ST10 strains. The proportion of each antigen and *fimH* allele has been plotted by year ordered by the overall frequency (most common at the bottom). Only the names of the five most frequent antigens and *fimH* alleles are shown. We did not detect significant variation for any of the five most frequent O and H groups nor for the five most frequent *fimH* alleles.

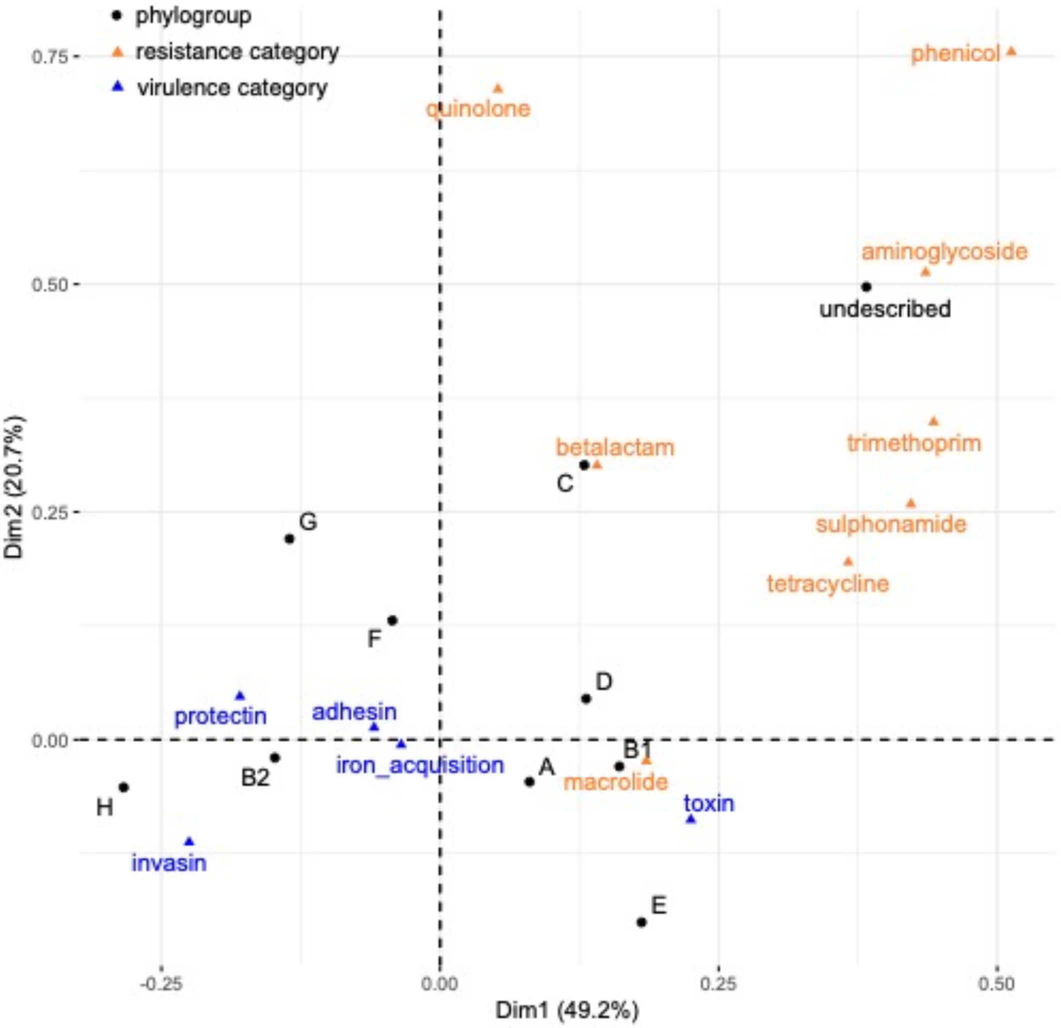


**Figure S8.** Projection of 21 variables, including virulence gene (in blue) and resistance categories (in orange), in the 403 isolates on the first two dimensions of the correspondence analysis (see table S11-13 for the contingency table, the eigenvalues and the contribution of rows and columns to the three main dimensions). For resistance, both gene acquisition and point mutations are included, but we omitted the polymyxin category because only one strain was resistant to polymyxin antibiotics. The phylogroup B2 contributes to 48% to the first axis (Dim1) and the phylogroups B1 and D to 16% and 13% to the first axis respectively. The phylogroups C, E and F contribute to 30%, 19% and 19% to the second axis (Dim2) respectively. The VFs (adhesin, invasin, iron acquisition and protectin categories) contribute to 28% to the first axis. The toxins contribute to 46% to the first axis and to 17% to the second axis. The large majority of resistance categories (resistances to beta-lactam, sulphonamide, aminoglycoside, phenicol, tetracycline and trimethoprim antibiotics) contribute to 21% to the first axis and to 52% to the second axis.

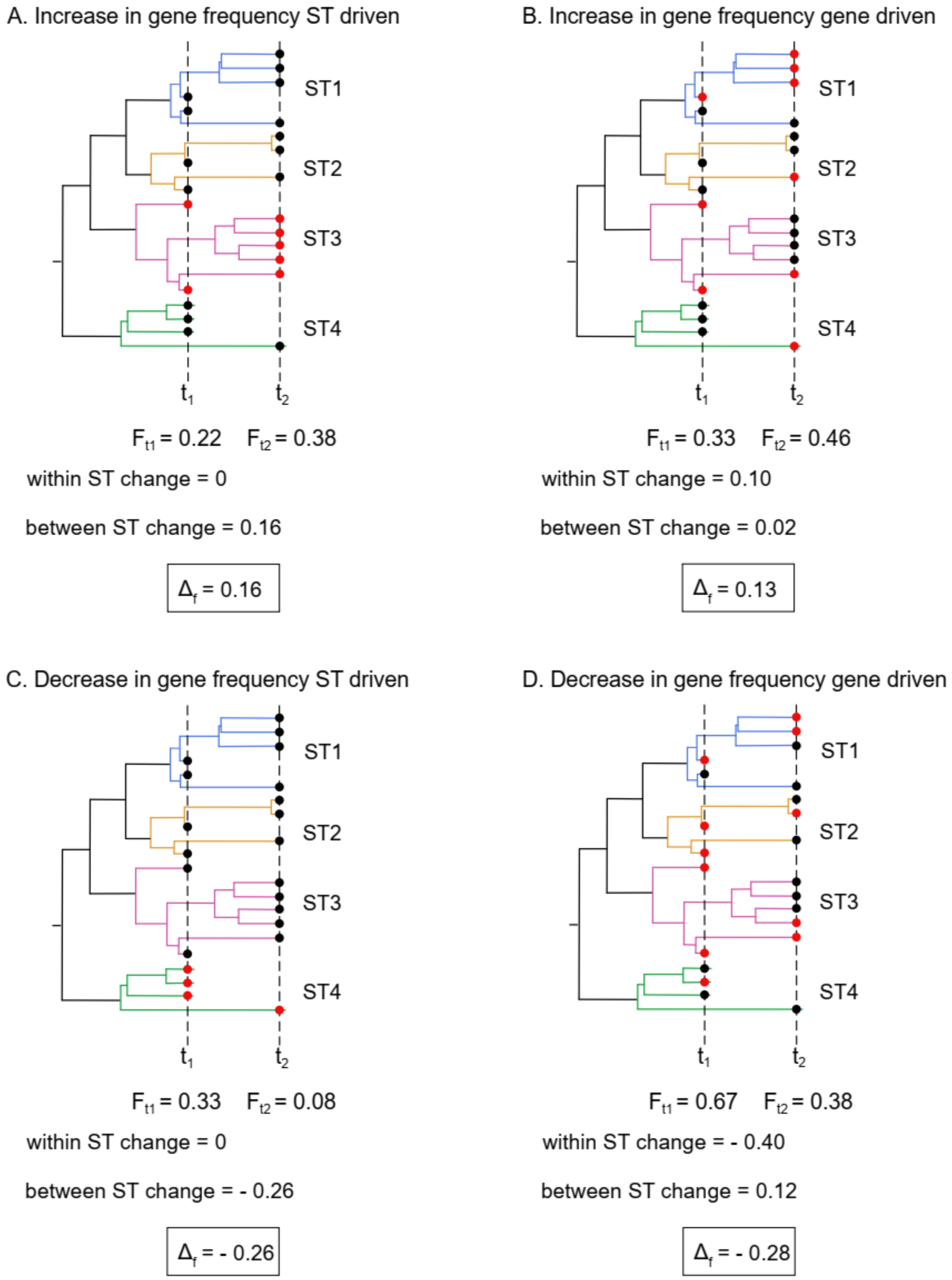


**Figure S9.** Decoupled temporal change in gene frequency. The overall change ($\Delta f$) is the sum of within ST change and between ST change. Nine strains were sampled at $t_1$ and 13 at $t_2$ (circles), with some of them carrying the focal gene (red circles).

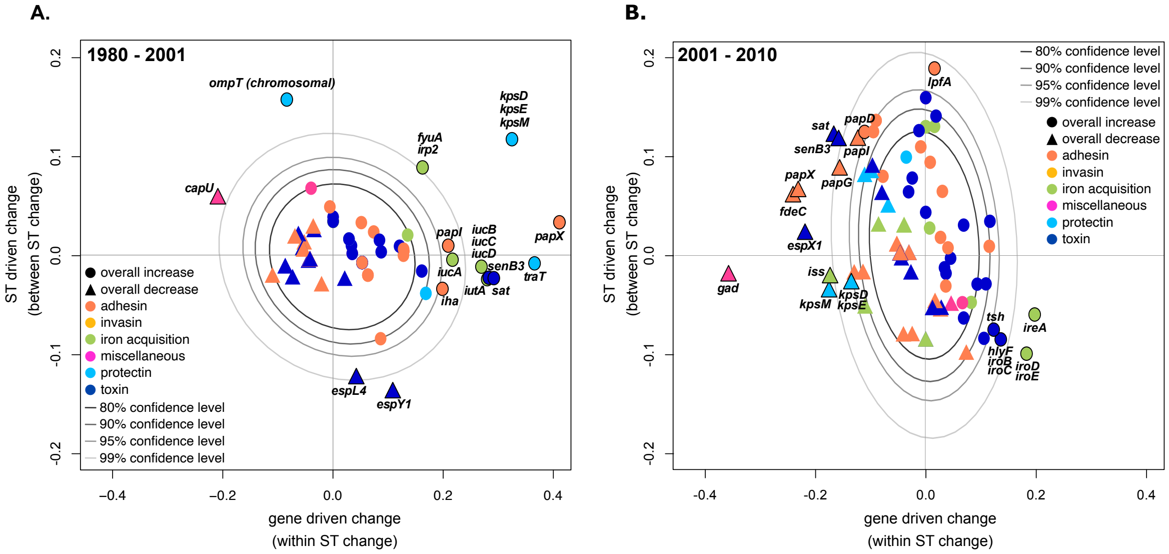


**Figure S10.** Temporal change of virulence gene frequency between 1980 – 2001 (A) and between 2001 – 2010 (B) excluding B2 strains. The overall frequency change ($\Delta f$) for each gene (increases depicted by circles and decreases by squares) is decomposed in two additive components, change driven by the variation in frequency of STs carrying the focal gene (ST driven change) and change driven by the variation in frequency of the focal gene (gene driven change). For readability, only genes for which between ST change or within ST change was greater than 0.02 in absolute value are shown (see table S17-18 for the complete list). The grey ellipses represent the confidence levels computed for NVNR genes. Genes highlighted in bold are those for which the temporal change is significantly different from NVNR genes at the 0.05 level (out of the 95% confidence level).

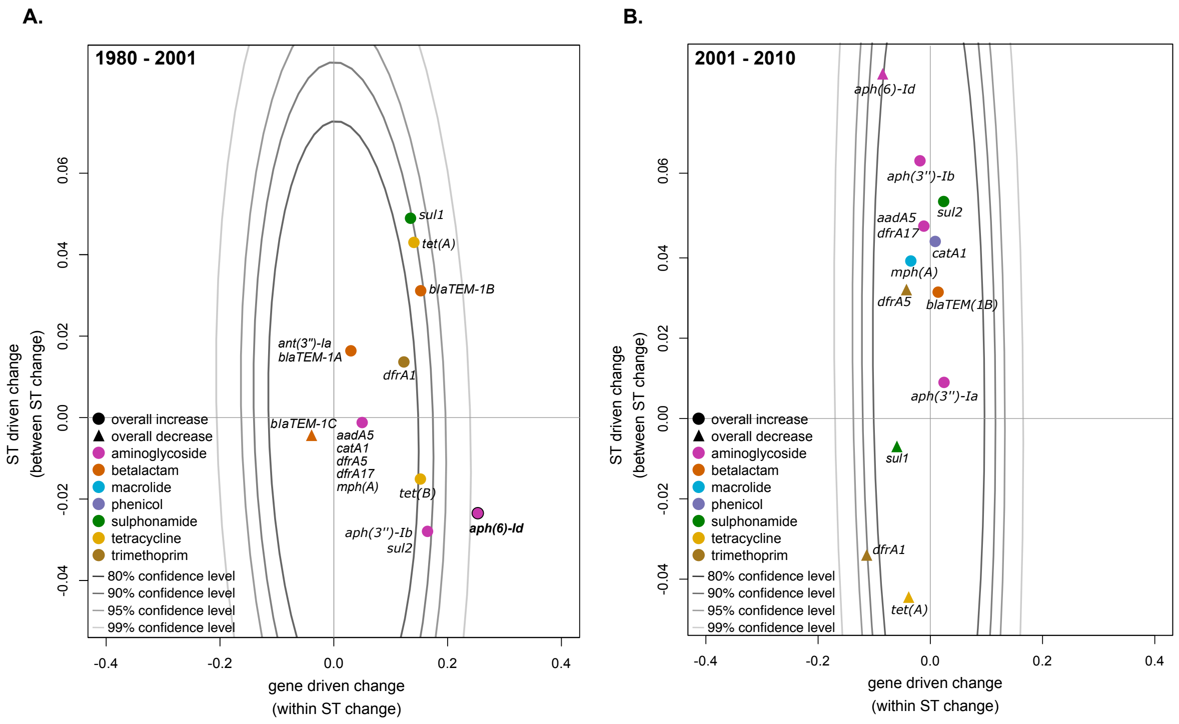


**Figure S11.** Temporal change of antibiotic resistance frequency between 1980 – 2001 (A) and between 2001 – 2010 (B) excluding B2 strains. The overall frequency change ($\Delta f$) for each gene (increases depicted by circles and decreases by squares) is decomposed in two additive components, change driven by the variation in frequency of STs carrying the focal gene (ST driven change) and change driven by the variation in frequency of the focal gene (gene driven change). For readability, only genes for which between ST change or within ST change was greater than 0.02 are shown here (see table S19-20 for the complete list). The grey ellipses represent the confidence levels computed for NVNR genes. Genes highlighted in bold are those for which the temporal change is significantly different from NVNR genes at the 0.05 level (out of the 95% confidence level). Note the scale of the y-axis is much smaller for resistance than for virulence genes (figure 4) as ST-driven changes are minor compared to gene-driven changes for antibiotic resistance genes.
